# Microbiota-instructed regulatory T cell differentiation is mediated by a distinct ROR*γ*t^+^ antigen presenting cell subset

**DOI:** 10.1101/2021.11.19.469318

**Authors:** Ranit Kedmi, Tariq Najar, Kailin R. Mesa, Allyssa Grayson, Lina Kroehling, Yuhan Hao, Stephanie Hao, Maria Pokrovskii, Mo Xu, Jhimmy Talbot, Jiaxi Wang, Joe Germino, Caleb A. Lareau, Ansuman T. Satpathy, Mark S. Anderson, Terri M. Laufer, Iannis Aifantis, Juliet M. Bartleson, Paul M. Allen, Helena Paidassi, James M. Gardner, Marlon Stoeckius, Dan R. Littman

## Abstract

The mutualistic relationship of gut-resident microbiota and cells of the host immune system promotes homeostasis that ensures maintenance of the microbial community and of a poised, but largely non-aggressive, immune cell compartment^1, 2^. Consequences of disturbing this balance, by environmental or genetic factors, include proximal inflammatory conditions, like Crohn’s disease, and systemic illnesses, both metabolic and autoimmune. One of the means by which this equilibrium is achieved is through induction of both effector and suppressor or regulatory arms of the adaptive immune system. In mice, *Helicobacter* species induce regulatory (iTreg) and follicular helper (Tfh) T cells in the colon-draining mesenteric lymph nodes under homeostatic conditions, but can instead induce inflammatory Th17 cells when iTreg cells are compromised^3, 4^. How *Helicobacter hepaticus* and other gut bacteria direct T cells to adopt distinct functions remains poorly understood. Here, we investigated which cells and molecular components are required to convey the microbial instruction for the iTreg differentiation program. We found that antigen presentation by cells expressing ROR*γ*t, rather than by classical dendritic cells, was both required and sufficient for iTreg induction. These ROR*γ*t^+^ cells, likely to be type 3 innate lymphoid cells (ILC3) and/or a recently-described population of Aire^+^ cells termed Janus cells^5^, require the MHC class II antigen presentation machinery, the chemokine receptor CCR7, and *α*v integrin, which activates TGF-β, for iTreg cell differentiation. In the absence of any of these, instead of iTreg cells there was expansion of microbiota-specific pathogenic Th17 cells, which were induced by other antigen presenting cells (APCs) that did not require CCR7. Thus, intestinal commensal microbes and their products target multiple APCs with pre-determined features suited to directing appropriate T cell differentiation programs, rather than a common APC that they endow with appropriate functions. Our results illustrate the ability of microbiota to exploit specialized functions of distinct innate immune system cells, targeting them to achieve the desired composition of equipoised T cells, thus maintaining tolerance.

## Main

A subset of bacterial species among the hundreds that comprise the gut microbiota elicit stereotypic antigen-specific T cell differentiation programs through mechanisms yet to be elucidated. It is generally accepted that conventional (or classical) dendritic cells (cDC) that migrate from tissue to the inductive sites in lymph nodes present microbial antigens to activate and promote differentiation of naive antigen-specific T cells^4, 6–10^. However, the antigen presenting cells (APCs) that execute these functions according to instructions of distinct intestinal bacterial species have not been clearly defined. We have chosen to study the APC requirements for T cell responses to *H. hepaticus* (Hh), which elicits iTreg, Tfh, or pathogenic Th17 cells under different conditions^3^.

### Selective requirement of APC subsets for gut microbiota-specific iTreg and inflammatory Th17 cell differentiation

To study the properties of antigen-presenting cells that direct the differentiation of microbiota- specific iTreg cells, we transferred into Hh-colonized mice naive Hh-specific CD4^+^ T cells from HH7-2 TCR transgenic mice^3^. CFSE-labeled transferred T cells exhibited robust proliferation by day 3 in colon-draining C1 mesenteric lymph nodes (MLN) of wild type (WT) mice, with up- regulation of Foxp3 and ROR*γ*t, characteristic of colonic iTreg cells^11, 12^. In contrast, in mice deficient for antigen presentation by DC (and, potentially, other cells: *CD11c-Cre;I-Ab^f/f^*, designated as *MHCII^ΔCD11c^)*, there was no expression of Foxp3 by the Hh-specific T cells, but, surprisingly, there was substantial proliferation of these cells, with up-regulation of ROR*γ*t (Fig. 1a). At 2-3 weeks after transfer, there was expansion in the colonic lamina propria of ROR*γ*t- and T-bet-expressing Hh-specific T cells in mutant mice, characteristic of a pro-inflammatory program (Extended Data Fig. 1a-c). Endogenous T cells also displayed this phenotype, with fewer Foxp3^+^ ROR*γ*t^+^ iTreg and expansion of Th17 cells (Extended Data Fig. 1b). This result suggested that antigen presentation by *Cd11c*-lineage cells is required for iTreg cell differentiation, but that it is dispensable for the differentiation of pro-inflammatory Th17 cells.

**Figure 1.**
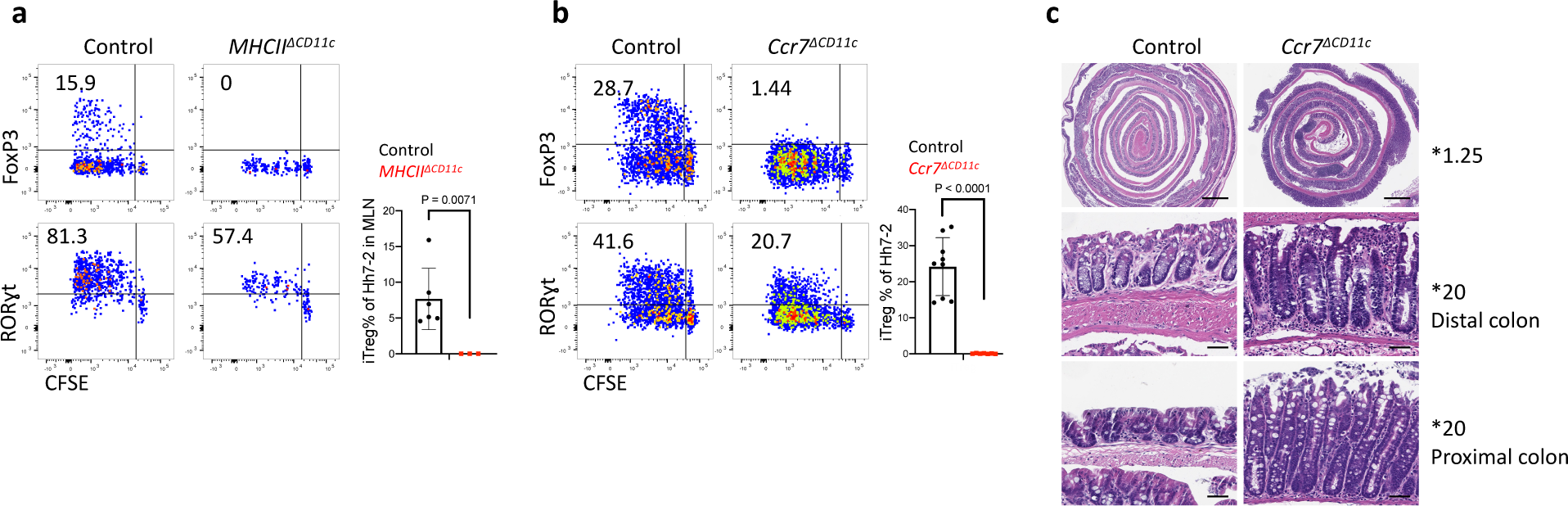
Distinct requirements for antigen presentation and CCR7 expression in differentiation of iTreg versus pathogenic Th17 cells. **a-b,** Hh-specific T cell proliferation and differentiation in Hh-colonized *CD11c-Cre;I-Ab^f/f^* (*MHCII^ΔCD11c^*) (n=3) and *I-Ab^f/f^* or *CD11c-Cre;I-Ab^f/+^* littermate control mice (n=6) (a) and in *Ccr7^Δ^*^CD11c^ (n = 7) and littermate control mice (n = 9) (b). CFSE-labeled naïve TCR transgenic Hh7-2 T cells from the C1 MLN were assessed for cell proliferation and expression of FoxP3 and ROR*γ*t at 3 days after adoptive transfer. Representative flow cytometry (left) and aggregate results (right). Data summarize two (a) and three (b) independent experiments. **c,** Representative H&E histology in large intestine of mice with indicated genotypes. Scale bars are 1 mm and 50µm, for 1.25X and 20X. respectively. All statistics were calculated by unpaired two-sided Welch’s t-test. Error bars denote mean ± s.d. *p-*values are indicated in the figure.

The chemokine receptor CCR7 mediates migration of DC and T cells into lymph nodes, where adaptive immune responses are initiated, and is critical for tolerogenic responses to food antigens^13^. In CCR7-deficient mice, iTreg induction in response to Hh colonization failed (Extended Data Fig. 1d), in agreement with the recent demonstration of a CCR7 requirement for the differentiation of iTreg cells specific for other *Helicobacter* species^6^. However, we observed robust priming and proliferation of ROR*γ*t^+^ Hh-specific T cells in CCR7-deficient mice (Extended Data Fig. 1d). In *CD11c-Cre;Ccr7^f/f^* conditional mutant mice^14^ (designated *Ccr7^ΔCD11c^*), transferred Hh-specific T cells failed to differentiate into iTreg cells, despite exhibiting robust proliferation, as in CCR7-deficient mice (Fig. 1b). In the colon of these mice, *Hh*-specific iTreg cells were rare, but there was accumulation of inflammatory Th17 cells along with elongation of the crypts (Fig. 1c, Extended Data Fig. 1e,f). Together, our results indicate that, unlike iTreg, microbiota-specific inflammatory Th17 cell differentiation does not depend on either CCR7 or antigen presentation by CD11c-expressing cells.

### Antigen presentation by *RORγt*-lineage cells is required for microbiota-induced iTreg cell differentiation

Classical DC, which have been broadly divided into cDC1 and cDC2, comprise multiple cell subsets that differ in their ontogeny, location, and transcription factor dependency^15, 16^. Both cDC1 and cDC2 have been proposed to initiate iTreg responses, largely based on their ability to induce Treg cell differentiation *in vitro*^9, 17^. However, *in vivo* depletion of cDC1 or cDC2 failed to phenocopy iTreg loss (Extended Data Fig. 2a-c), in agreement with previous reports^6, 18–20^. These findings suggested that antigen presentation by a rare uncharacterized *Cd11c*-lineage myeloid or non-myeloid cell subset is required for iTreg cell differentiation. To identify putative antigen presenting cell populations targeted by *CD11c-cre*, we performed CITE-seq analysis of cells isolated from MLN of Hh-colonized *CD11c-Cre;ROSA26LSLtdTomato* (designated *tdTomato- ON^ΔCD11c^*) fate-map mice (Fig. 2a and Extended Data Fig. 3a,b). In addition to expected myeloid cell subsets, we identified both ILC3 and a recently described Aire^+^ ROR*γ*t^+^ population, named Janus cells (JC)^5, 21^, among the tdTomato^+^ cells, and these also expressed MHCII (Fig. 2b and Extended Data Fig. 3b). Using a gating strategy for JC (Extended Data Fig. 4a-d), we confirmed that they, like ILC3, express *Rorc* (Extended Data Fig. 3a). Consistent with this, GFP^+^ ILC3 and JC from the C1 MLN of *RORγt-eGFP* mice expressed both CD11c and CD11b or mostly CD11c, respectively and were both targeted by *CD11c-Cre* (Fig. 2c, Extended Data Fig. 3b and 4e). Accordingly, many fewer ROR*γ*t^+^ cells from MLN of *MHCII^ΔCD11c^* mice expressed MHCII, as compared to WT littermates (Extended Data Fig. 5a).

**Figure 2.**
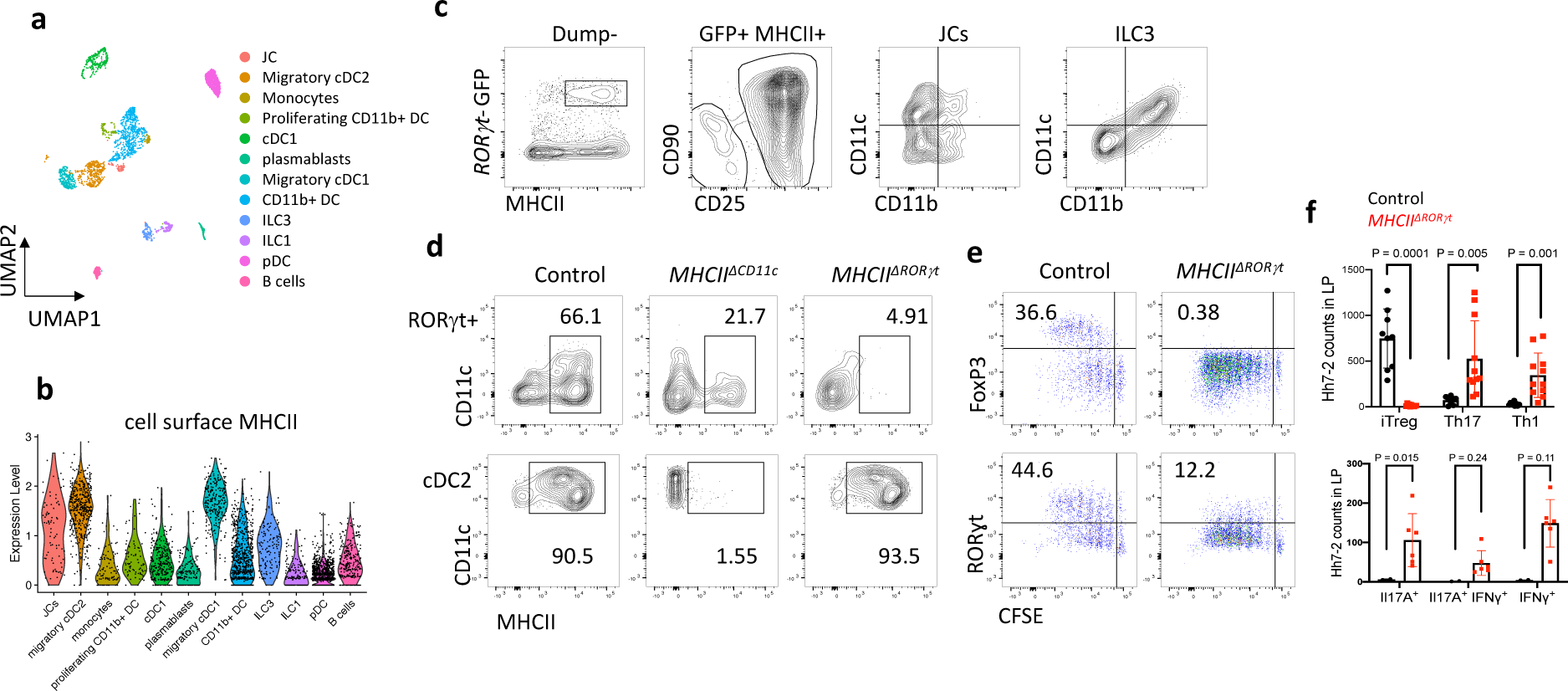
Antigen presentation by ROR*γ*t^+^ cells is required for microbiota-induced iTreg cell differentiation. **a-b,** UMAP visualization of the *tdTomato-ON^CD11c^* fate-map cell CITE-seq dataset, analyzed by the WNN method (a), and Violin plot showing MHCII protein levels in the different cell clusters (b). MLN cells from Hh-colonized *tdTomato-ON^CD11c^* fate-map mice were gated for TCR*β*^-^, TCR*γδ*^-^, B220^-^, and tdTomato^+^ cells were sorted for CITE-seq analysis. Cells were sorted from two mice and labeled by hashing antibodies (n=2). **c,** CD11c and CD11b staining of ILC3 and JC from MLN of Hh-colonized *RORγt-eGFP* mice, gated as indicated. **d,** MHCII expression in ROR***γ***t^+^ cells (top) and DCs (bottom) from the MLN of Hh-colonized mice of the indicated genotypes. ROR***γ***t^+^ cells were gated as TCR*β*^-^, TCR*γδ*^-^, B220^-^, ROR*γ*t^+^; DC were gated as TCR*β*^-^, TCR*γδ*^-^, B220^-^, CD90^-^, CD11c^+^, CD11b^+^ SIRP*α*^+^. **e,** Hh7-2 cell proliferation and differentiation in *MHCII^ΔRORγt^* (n = 6) and *I-Ab^f/f^* littermate control mice (n = 6) at 3 days after adoptive transfer into Hh-colonized mice. **f,** Hh7-2 T cell differentiation profiles (upper) and cytokine production (lower) in the large intestine lamina propria at 22 days after transfer into *MHCII^ΔRORγt^* (n=11) and littermate controls (n=9). Differentiation was assessed by expression of Foxp3, ROR*γ*t with or without T-bet, and T-bet. Data summarize two independent experiments. All statistics were calculated by unpaired two-sided Welch’s t-test. Error bars denote mean ± s.d. *p*-values are indicated on the figure.

**Figure 3.**
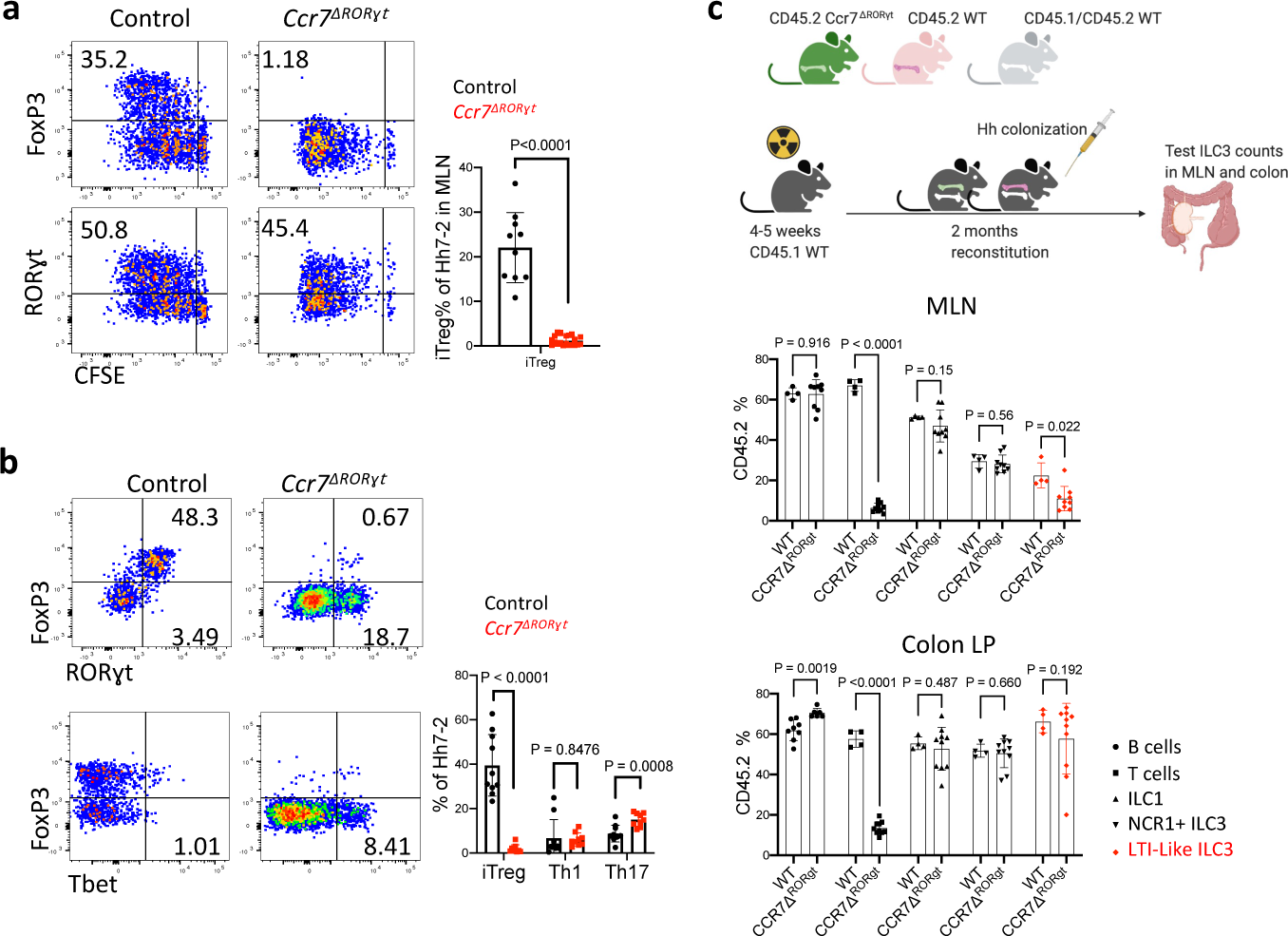
ROR*γ*t^+^ cells require CCR7 to promote iTreg cell differentiation. **a-b,** Representative flow cytometry profiles (left) and aggregate data (right) for Hh7-2 T cell proliferation and differentiation in MLN at 3 days (a) and their phenotype in large intestine at 14 days (b) following adoptive transfer into *Ccr7^ΔRORγt^* and littermate control mice. MLN: Control mice n=10, *Ccr7^ΔRORγt^* mice n=20; LI: Control mice n=10, *Ccr7^ΔRORγt^* mice n=9. Data summarize three independent experiments. Statistics were calculated by unpaired two-sided Welch’s t-test. **c,** Analysis of CCR7 requirement for ROR*γ*t^+^ cell accumulation in the MLN. Irradiated CD45.1 mice were reconstituted with equal number of bone marrow cells from CD45.2 *Ccr7^ΔRORγt^* and CD45.1/CD45.2 WT mice or with CD45.2 WT and CD45.1/CD45.2 WT mice as controls. Scheme is shown at the top (created with BioRender.com). Aggregate data shows the frequency in MLN and colon lamina propria of CD45.2 WT (n=4) or CD45.2 *Ccr7^ΔRORγt^* (n=9) cells within each subset, as indicated. Data summarize three independent experiments. All statistics were calculated by unpaired two-sided Welch’s t-test. Error bars denote mean ± s.d. *p*- values are indicated in the figure.

**Figure 4.**
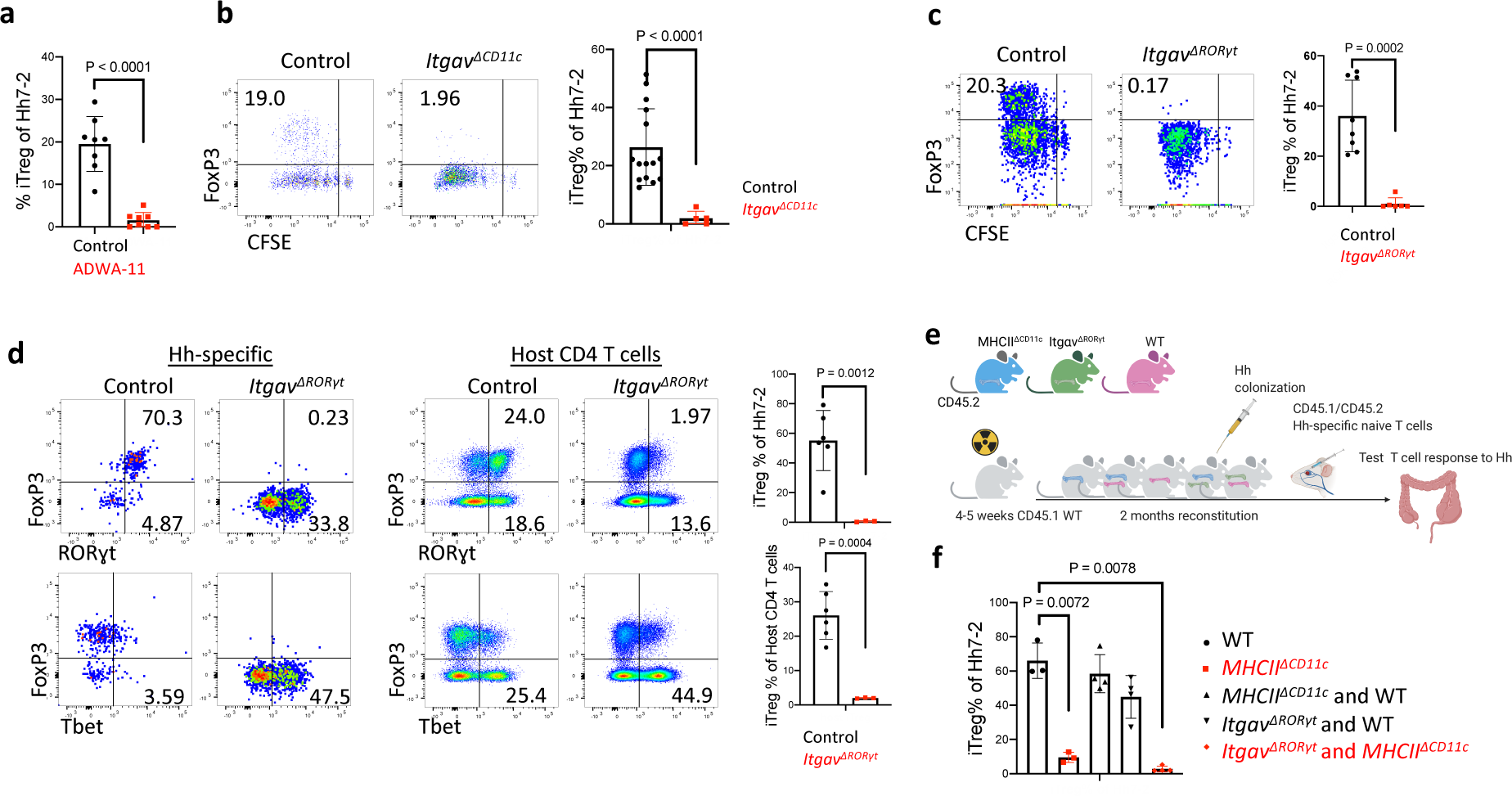
Role of integrin *α*v*β*8 in ROR*γ*t^+^ antigen presenting cell-dependent iTreg cell differentiation. **a,** Frequency of iTreg cells among proliferating donor-derived Hh-specific cells in the MLN at 3 days after transfer of naïve CFSE-labeled Hh7-2 T cells into mice treated with 200μg of ADWA11 blocking antibody (n=8) or left untreated (n=8), on the day of adoptive transfer. Data summarize three independent experiments. **b,** Hh7-2 T cell proliferation and differentiation in the MLN of *Itgav^ΔCD11c^* (n=5) and littermate controls (n=15) at 3 days after adoptive transfer. Data summarize three independent experiments. **c,** Proliferation and differentiation of Hh-specific iTreg cells in the MLN of *Itgav^ΔRORγt^* (n=6) and littermate control mice (n=8). CFSE-labeled Hh7-2 T cells were analyzed at 3 days following their adoptive transfer into Hh-colonized mice. Representative flow cytometry profiles (left) and aggregate data (right). Data summarize three independent experiments. **d,** Transcription factor expression in Hh7-2 T cells (left panels) and in endogenous CD4^+^ T cells (right panels) from colon lamina propria (LP) at 10 days after adoptive transfer into *Itgav^ΔRORγt^* mice (n=3) and control littermates (n=6). Data summarize two independent experiments. Representative dot plots and aggregate data are shown (right panels). **e,** Scheme for mixed bone marrow chimeric mouse experiment, with control, *Itgav^ΔRORγt^* or *MHCII^ΔCD11c^* cells administered to irradiated host mice (created with BioRender.com). **f,** Bar graphs showing iTreg frequency among Hh7-2 T cells in the colon LP at 10 days after their transfer into the bone marrow chimeric mice, reconstituted with different combinations of donor cells as indicated. Control mice (n=3), *MHCII^ΔCD11c^* (n=3), *MHCII^ΔCD11c^* and WT (n=4), *Itgav^ΔRORγt^* and WT (n=4) and *MHCII^ΔCD11c^* and *Itgav^ΔRORγt^* (n=4). All statistics were calculated by unpaired two-sided Welch’s t-test. Error bars denote mean ± s.d. *p-*values are indicated in the figure.

**Figure 5.**
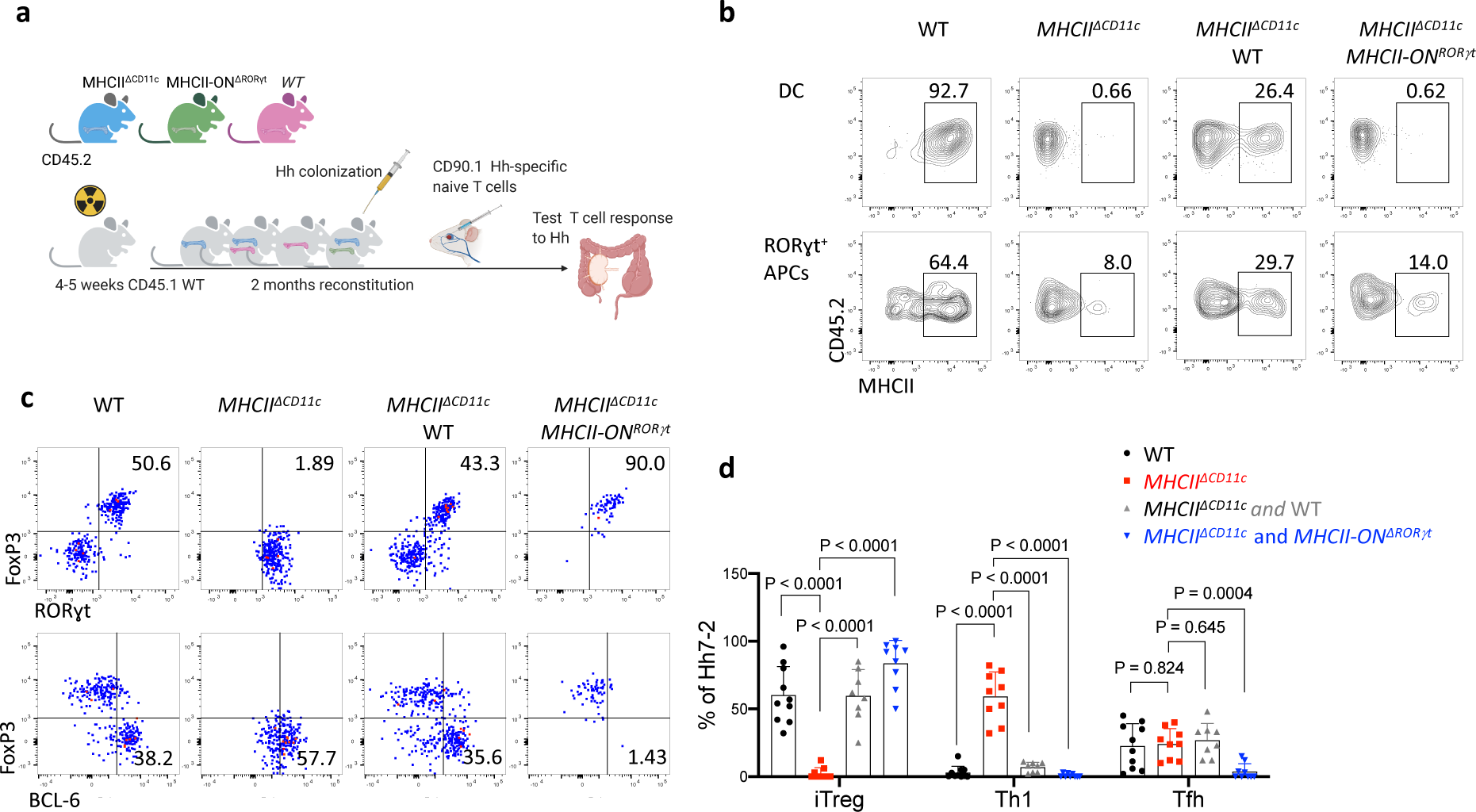
Antigen presentation by ROR*γ*t^+^ cells is sufficient to promote iTreg cell differentiation. **a,** Experimental design (created with BioRender.com). **b,** MHCII frequency in donor bone marrow-derived cDC2 (gated TCR*β*^-^, TCR*γδ*, B220^-^, CD45.2^+^, CD11c^+^, CD11b^+^ Sirpa^+^) and ROR*γ*t^+^ cells (gated as TCR*β*^-^, TCR*γδ*^-^, B220^-^, RORγt^+^, CD45.2^+^) in MLN from chimeric mice reconstituted with combinations of donor BM cells as indicated. **c,** Representative flow cytometry of Hh7-2 T cell differentiation in colon lamina propria of Hh-colonized bone marrow chimeric mice, 12 days after transfer of naive TCR transgenic T cells. **d,** Aggregate data for differentiation of Hh7-2 T cells in bone marrow chimeric mice reconstituted with cells of indicated genotypes. WT (n=10), MHCII*^ΔCD11c^* (n=9), MHCII*^ΔCD11c^* and WT (n=8), MHCII*^Δ^*^CD11c^ and MHCII-ON^ΔRORγt^ (n=9). Data summarize three independent experiments. All statistics were calculated by unpaired two-sided Welch’s t-test. Error bars denote mean ± s.d. *p-*values are indicated in the figure.

Expression of MHCII by ILC3 was reported to prevent microbiota-dependent expansion of inflammatory T cells in the intestine, and was proposed to mediate negative selection of those cells^22^. Our results suggested that *CD11c-Cre*-expressing ROR*γ*t^+^ cells may, instead, be required for the differentiation of microbiota-specific iTreg cells. This was confirmed by examining iTreg cell differentiation in mice whose MHCII was inactivated in presumptive ILC3 and JC (*RORγt- Cre;I-AB^l/l^,* designated as *MHCII^ΔRORγt^*). Despite expression of MHCII in cDC2 of these mutant mice (Fig. 2d), there was complete loss of Hh-specific iTreg cell differentiation in colon-draining MLN, but intact priming and subsequent expansion of pathogenic Th17 cells in the large intestine lamina propria (Fig. 2e,f, Extended Data Fig. 5b). Both the donor Hh-specific and host T cells exhibited loss of iTreg and increase in IFN*γ*- and IL-17A-producing CD4^+^ T cells in the large intestine lamina propria (Fig. 2f, Extended Data Fig. 5c-e). Similar results were observed with Hh-specific T cells in another conditional mutant strain, *RORγt-Cre; H2-DMa^f/f^* mice (*H2-DMa^ΔRORγt^*) deficient in H2-DMa, the mouse equivalent of HLA-DM, required for displacement of invariant chain peptide and loading of processed peptide on MHCII molecules^23^. This result confirms that the antigen processing machinery is required in ROR*γ*t^+^ APCs for induction of microbiota-specific iTregs (Extended Data Fig. 5f).

### CCR7-dependent migration by *RORγt*-lineage cells is required for microbiota-induced iTreg cell differentiation

Intestinal ILC3 have been reported to also employ CCR7 for migration to draining MLN^24, 25^. We found expression of *Ccr7* in both ILC3 and JC (Extended Data Fig. 3a and 6a). We therefore asked whether CCR7 is required for migration of ROR*γ*t ^+^ APCs rather than classical DC for iTreg induction. Indeed, Hh-specific iTreg cell differentiation was abrogated in the MLN of *Ccr7^ΔRORγt^* mice (Fig. 3a). In the colon of *Ccr7^ΔRORγt^* mutant mice, there was skewing of Hh-specific T cells towards a Th1-Th17 inflammatory program after 2-3 weeks of adoptive T cell transfer, although this was less marked than in *Ccr7^ΔCD11c^* mice, and elongation of crypts was not observed (Fig. 3b and data not shown). To determine whether loss of CCR7 affects migration of ROR*γ*t^+^ cells to the MLN, we reconstituted irradiated mice with equal numbers of WT and *RORγt-Cre;Ccr7^f/f^* (*Ccr7^ΔRORγt^*) bone marrow cells. Although there was no effect on the ratio of WT and mutant migratory DC, there was a substantial reduction in the proportion of CCR7-deficient lymphoid tissue inducer (LTi)-like ILC3, but no significant difference in the proportions of other ROR*γ*t^+^ cells in the MLN (Fig 3c and Extended Data Fig. 6b-d). Similarly, in *Ccr7^ΔCD11c^* mice, LTi-like ILC3 lost CCR7 expression and were reduced in number in the MLN (Extended Data Fig. 6e,f). To further study a potential role for JC, we examined the T cell response to Hh in *RORγt-cre;Aire^f/f^* mice and in Aire-DTR bone marrow-reconstituted mice treated with diphtheria toxin. In neither case was there an effect on Hh-specific iTreg cell differentiation (Extended Data Fig. 7a,b). We cannot, however, rule out a role for JC, as Aire may not be required for the microbiota-dependent iTreg inductive function of these cells and residual cells in DT-treated animals may be sufficient to support iTreg cell differentiation. Together, these results indicate that ROR*γ*t^+^ APC, either LTi-like ILC3 or JC, migrate to the MLN, where they present microbial antigen to naïve T cells to induce iTreg cell differentiation, and that their failure to migrate results, instead, in inflammatory Th17 cell differentiation.

### Integrin *α*v expression by *RORγt*-lineage antigen presenting cells is required for iTreg differentiation

Differentiation of iTreg cells requires TGF-β signaling in CD4^+^ T cells, and defects in this pathway result in spontaneous colitis^3, 11^. TGF-β is released from its latent form on cell surfaces or extracellular matrix following physical interaction with integrins *α*vβ6 or *α*vβ ^26–28^. Loss of integrins β8 or *α*v in hematopoietic cells, including in *CD11c-Cre:Itgb8^f/f^* mice, resulted in reduced colonic Tregs and in multiorgan inflammation^29–31^. Consistent with those observations, differentiation of adoptively transferred Hh-specific T cells into iTreg cells was abrogated both following treatment of mice with anti-β8 antibody and in *CD11c-Cre;Itgαv^f/f^* (Itgav*^ΔCD11c^*) recipient mice (Fig. 4a,b). We examined iTreg cell differentiation in mice with conditional inactivation of *Itgαv* in ROR*γ*t^+^ APCs and in T cells (*Itgav^ΔRORγt^*) (Fig. 4c,d). In the colon-draining MLN of *Itgav^ΔRORγ^* mice, there was loss of integrin *α*v (CD51) expression on both ILC3 and JC, and adoptively transferred Hh-specific T cells failed to express Foxp3 (Fig. 4c and Extended Data Fig. 8a). The MLN were increased in size (Extended Data Fig. 8b) and the Hh-specific T cells expressed ROR*γ*t, but, unlike in control littermates, they also had elevated T-bet, along with a substantial decrease in CCR6, consistent with reduced TGF-β signaling (Extended Data Fig. 8c). Notably, β8 antibody blockade resulted in the same phenotype (Extended Data Fig. 8d). In the colonic lamina propria of *Itgav^ΔRORγt^* mice, there was loss of both Hh-specific and host-derived iTreg cells, with skewing of CD4^+^ T cells towards IFN*γ*^+^ Th1 and pathogenic Th17 programs (Fig. 4d, Extended Data Fig. 8e), suggesting that expression of integrin *α*v on ROR*γ*t+ APCs, likely in partnership with integrin β8, is a general requirement for intestinal iTreg cell differentiation. Although ILC3 express higher levels of CD51 (Extended Data Fig. 8a), single-cell RNA sequencing analysis of GFP^+^ cells from pooled lymph nodes of *Aire* reporter mice^32^ showed clustering of JCs into three discrete subpopulations, with JC2 and JC3 expressing high levels of *Itgav* and *Itgb8* (Extended Data Fig.9a-e). JC and a fraction of ILC3, obtained from *Itgb8-tdTomato* reporter mice^33^, were found to express tdTomato (Extended Data Fig 9f), and we therefore cannot exclude the requirement of either cell type for iTreg differentiation. Differentiation of iTreg cells was normal in *CD4-Cre;Itgαv^f/f^* mice (Extended Data Fig. 8f), consistent with a role of the integrin in ROR*γ*t^+^ APCs rather than in TCR*αβ* T cells. Furthermore, we reconstituted mice after irradiation with a mix of MHCII*^ΔCD11c^* and *Itgav^ΔRORγt^* bone marrow cells, resulting in binary expression of MHCII or integrin *α*v. In these mice, iTreg cell differentiation was abolished, consistent with a requirement for both antigen presentation and activation of TGF-β by the same APC, coupling T cell activation with differentiation cues (Fig. 4e- f, Extended Data Fig. 8g).

### Antigen presentation by ROR*γ*t^+^ APCs is sufficient to promote iTreg cell differentiation

Our results support a role for *RORγt^+^* APCs in microbiota-specific iTreg differentiation, but do not rule out a requirement for additional conventional antigen presenting cells. Moreover, a population of DC (T-bet-negative cDC2) was recently reported to express ROR*γ*t^34^, raising the possibility that neither ILC3 nor JC is relevant in iTreg cell differentiation. To examine a potential role for a rare DC subset, we employed *zbtb46-Cre*, considered to specifically target cDC, and *zbtb46* reporter mice. Because *zbtb46-Cre;Ccr7^f/-^* mice (CCR7^Δzbtb46^) were unable to support microbiota- dependent iTreg cell differentiation (Extended Data Fig. 10a), we profiled by CITE-seq sorted cells from C1 mLN of *zbtb46-eGFP ; RORγt-Cre;ROSA26LSLtdTomato* (*tdTomato-ON^RORγt^*) mice, expressing one or both fluorescent reporters, gated to exclude B and T cells (Extended Data Fig. 10b). Surprisingly, GFP expression was identified on all ILC3 and fate-mapped JC. Zbtb46 expression on ILC3 was confirmed using *mKate2-ON^zbtb^*^46^; *RORγt-eGFP* mice (Extended Data Fig. 10c). We identified a few migratory cDC2 among tdTomato^+^ GFP^+^ cells in *zbtb46-eGFP; tdTomato-ON^RORγt^* mice, but these did not form a subcluster to suggest a unique gene signature and did not exhibit active *Rorc* or integrin *α*v mRNA and protein expression (Extended Data Fig. 10d-f), suggesting that they are unlikely to be ROR*γ*t^+^ APCs required to direct iTreg cell differentiation. We performed three-dimensional intravital imaging studies to visualize interactions of newly-primed Hh-specific T cells with DC and *RORγt^+^* APCs populations in C1 MLN. We utilized *mKate2-ON^zbtb46^*; *RORγt-eGFP* mice to visualize cDC (mKate2^+^) and *RORγt^+^* APCs (eGFP^+^ and mKate2). However, given that the efficiency of *zbtb46-Cre*-mediated activation of the mKate2 reporter was low (∼20% for *RORγt+* APCs and cDC populations) (Extended Data Fig. 11a), we additionally utilized cell morphology and size analysis to distinguish the few host-derived GFP^+^ T cells from GFP^+^ *RORγt^+^* APCs. We next transferred dye-labeled *Nur77-eGFP* Hh-specific T cells into these fluorescent reporter host mice to measure direct interactions of primed T cells (which up-regulate *Nur77*) with DC and *RORγt^+^* APCs at 15 h after adoptive transfer. Approximately 81% of GFP^+^ primed Hh-specific T cells were found in contact with at least one *RORγt^+^* APC with/without DC, as opposed to only ∼31% of the non-primed T cells (Extended Data Fig. 11b,c).

Our imaging study and results with conditional mutant mice did not rule out a contribution by DC towards Hh-specific T cell activation and iTreg differentiation. We therefore wished to determine whether antigen presentation limited to only *RORγt-Cre*-expressing cells was sufficient to allow for iTreg cell differentiation. For this purpose, we used mice that express MHCII only in *RORγt^+^* APCs, and not in DC or other APCs (*RORγt-Cre;I-AB^-/lsl^*, designated *MHCII-ON^RORγt^*). We reconstituted irradiated congenic mice with bone marrow from *MHCII^ΔCD11c^* mice with or without bone marrow from WT or *MHCII-ON^RORγt^* mice (Fig. 5a). Flow cytometry analysis confirmed MHCII expression by *RORγt^+^* cells, but not DC from the C1 MLN and the colon lamina propria of *MHCII^ΔCD11c^*;*MHCII-ON^RORγt^* bone marrow-reconstituted mice (Fig. 5b, Extended Data Fig. 12a). As expected, in control mice reconstituted with only *MHCII^ΔCD11c^* bone marrow cells, in which no MHCII expression was detected in either *RORγt^+^* APC or DC, there was no differentiation of adoptively transferred Hh-specific iTreg cells, but there was, instead, differentiation of ROR*γ*t^+^Tbet^+^ inflammatory Th17 cells (Fig. 5c,d). In contrast, antigen presentation by ROR*γ*t^+^ cells alone was sufficient to rescue iTreg cell differentiation and suppression of inflammatory T cells in response to Hh colonization, as seen in mice reconstituted with *MHCII^ΔCD11c^* plus *MHCII-ON^RORγt^* bone marrow cells (Fig. 5c,d), or with only *MHCII-ON^RORγt^* cells (Extended Data Fig. 12c). There was similar rescue of endogenous iTreg cell differentiation in mice having the gain-of-function MHCII in ROR*γ*t^+^ cells (Extended Data Fig. 12b). It should be noted that whereas there was rescue of Hh-directed iTreg cell differentiation in mice reconstituted with *MHCII-ON^RORγt^* bone marrow, there was a marked absence of Bcl6-expressing Tfh cells, which are also induced by Hh (Fig. 5c,d; Extended Data Fig. 12c). Interestingly, Tfh cells were present in mice reconstituted with only *MHCII-ON^CD11c^* bone marrow, suggesting that their differentiation requires antigen presentation by DC and/or B cells, as proposed previously^35^ (Extended Data Fig. 12c). We conclude that *RORγt^+^* APC, ILC3 and/or JC, are specialized to prime naive microbiota-specific T cells and guide their differentiation into iTregs, but other APCs are required to guide the differentiation of microbiota-specific pathogenic Th17 cells and Tfh cells.

## Discussion

The composition of the intestinal microbiota influences host immune functions that contribute to anti-microbial host defense, inflammatory disease, and anti-tumor immunity. Transmission of information from gut microbes to immune system cells remains poorly understood. The current results indicate that ROR*γ*t^+^ cells, either JC, whose transcriptional profile suggests a role in promoting immunological tolerance^5^, or type 3 innate lymphoid cells previously implicated in restraining microbiota-dependent Th1/Th17 inflammatory responses in the gut^36, 37^, do so in large part by conveying signals from the microbiota to naive bacteria-specific T cells, activating them and guiding their differentiation towards a unique iTreg cell program. ROR*γ*t^+^ cells defective for CCR7-mediated migration (either to or within the MLN), MHCII antigen presentation, or αvβ8 function (presumably through activation of TGF-β), failed to induce the iTreg program. Intriguingly, under such circumstances there was expansion of pathogenic Th17 cells that promoted intestinal inflammation. We previously demonstrated that a T cell-intrinsic c-Maf deficiency prevented iTreg cell differentiation, and similarly allowed for microbiota-dependent differentiation of pathogenic Th17 cells^3^. Together, these findings suggest that iTreg cells restrain the priming, proliferation, and differentiation of Th17 cells in the MLN. The APC(s) that directs pathogenic Th17 cell differentiation does not require CCR7, and it is not targeted in CD11c-Cre mice, and is, therefore, most likely not a cDC, but its identity is currently not known. One hypothesis is that iTreg cells might not only inhibit Th17 cell differentiation and function but also may inhibit the function of the CCR7-independent Th17 inducer APC (Extended Data Fig. 13).

It has been proposed that intrinsically different APC subsets direct distinct T cell responses^38^, but such processes have been difficult to demonstrate in the setting of immune responses *in vivo*. Our study shows that a unique RORγt^+^ cell type instructs naïve microbiota-specific CD4^+^ T cells to become iTreg cells, but does not support the differentiation of other T cell programs, including Tfh cells, that are normally also induced by Hh intestinal colonization. Although our results are most compatible with the ROR*γ*t^+^ APC being an ILC3 or Aire^+^ JC subset, we cannot rule out that it may represent a novel ROR*γ*t^+^ cell type that cannot yet be categorized as either lymphoid or DC-like. Definitive identification and characterization of this cell awaits more specific genetic tools than those currently available. Nevertheless, our results clearly demonstrate the existence of multiple APCs that are targeted by a specific commensal microbe to instruct diverse effector T cell functions (Extended Data Fig. 13). The APCs may act hierarchically, as exemplified here by ROR*γ*t^+^ cells that supersede the function of Th17-inducing APC. The existence and identity of distinct cellular circuits responsible for the induction of iTregs and other T cell functional subsets offers the opportunity to investigate the corresponding cells in humans and, potentially, to modulate them therapeutically.

## Acknowledgements

We thank members of the Littman lab, Juan J. Lafaille, and Susan Schwab for valuable discussion and critical reading of the manuscript and Gabriela Romero-Meza for assistance with experiments. We thank Dean Sheppard for advice and providing ADWA11 blocking Ab, S.Y. Kim and the NYU Rodent Genetic Engineering Laboratory (RGEL) for rederivation of mutant mice, and Cindy Loomis and the Experimental Pathology Research Laboratory of NYULMC for histology of intestine samples. The Microscopy Core and the Genome Technology Core are partially supported by NYU Cancer Center Support Grant NIH/NCI P30CA016087 at the Laura and Isaac Perlmutter Cancer Center, S10 RR023704-01A1 and NIH S10 ODO019974-01A1. The Experimental Pathology Research Laboratory is supported by National Institutes of Health Shared Instrumentation grants S10OD010584-01A1 and S10OD018338-01. Caleb A. Lareau, Ansuman T. Satpathy and James M. Gardner are recipients of the IGVF award UM1HG012076. This work was supported by an Irvington Institute fellowship from the Cancer Research Institute (R.K.) and a Jane Coffin Childs Fund fellowship (K.R.M.), the Helen and Martin Kimmel Center for Biology and Medicine (D.R.L.); National Institutes of Health grants R01AI139540 (P.M.A.) and R01AI158687 (D.R.L.), and the Howard Hughes Medical Institute (D.R.L.).

## Author Contributions

R.K., T.N., K.R.M. and D.R.L. designed the study and analyzed the data; R.K. and T.N. performed mouse genetic experiments with assistance from A.G; M.P., M.X., and J.T. performed early experiments to launch the study. Intravital multiphoton microscopy (K.R.M. and R.K.), CITE-seq studies (R.K, S.H. and M.S.), scRNA-seq (A.T.S., C.A.L), bioinformatics analyses (R.K, L.K., Y.H., J.G.). J.W., M.S.A., and J.M.G. provided biological samples, genomics data, and experimental support. H.P., T.M.L., I.A., J.M.B., and P.M.A. contributed mouse strains and phenotypic analysis (H.P.). R.K. and D.R.L. wrote the manuscript, with input from the other authors. D.R.L. supervised the research.

## Methods

### Mice

Mice were bred and maintained in the Alexandria Center for the Life Sciences animal facility of the New York University School of Medicine, in specific pathogen-free conditions. C57BL/6 mice (Jax# 000664), *Batf3^-/-^(B6.129S(C)-Batf3tm1Kmm/J* #Jax 013755), *Itgav^f/f^* (*B6.129P2(Cg)- Itgav^tm2Hyn^/J* Jax# 032297, CD45.1 mice (*B6.SJL-Ptprca Pepcb/BoyJ*, Jax# 002014), *CD4-Cre* (*Tg(Cd4-cre)1Cwi/BfluJ,* Jax# 017336), *CD11c-Cre* (*B6.Cg-Tg(Itgax-cre)1-1Reiz/J* #Jax 008068), *Ccr7^-/-^*(*B6.129P2(C)-Ccr7tm1Rfor/J*, Jax# 006621), *I-AB^f/f^* (*B6.129X1-H2-Ab1tm1Koni/J* #Jax 013181), *Zbtb46-Cre* (*B6.Cg-Zbtb46tm3.1(cre)Mnz/J* #Jax 028538), *Zbtb46-eGFP* (*B6.129S6(C)-Zbtb46tm1.1Kmm/J* #Jax 027618), tdTomato^LSL^ (*B6;129S6-Gt(ROSA)26Sortm14(CAG-tdTomato)Hze/J* #Jax 007908), *Aire^f/f^* (*B6.Cg-Airetm1Dfil/J* #Jax 031409), *Nur77-eGFP* (*C57BL/6-Tg(Nr4a1-EGFP/cre)820Khog/J* #Jax 016617), CD90.1 (*B6.PL-Thy1a/CyJ* #Jax 000406) mice were purchased from Jackson Laboratories. RORγt-Cre and Hh7-2tg were generated in our laboratory and previously described^3, 39^. BAC-transgenic Rorc(t)-GfpTG were generated by G. Eberl’s lab^40^. *huLangerin (CD207)-DTA* mice were kindky provided by Daniel H. Kaplan^41^. Ccr7*^f/f^* mice were previously described^14^. *mKate2^LSL^* mice^42^ were provided by Scott Lowe. I-AB lox-STOP-lox^35^, Aire-DTR^43^, H2-DMa1*^f/f^* ^23^, *Itgb8-tdTomato*^33^ Adig (Aire-Driven Igrp-Gfp)^32^ mice have been described. Littermates with matched sex (both males and females) were used. Mice in all the experiments were 6–12 weeks old at the starting point of treatment. Animal sample size estimates were determined using power analysis (power=90% and alpha=0.05) based on the mean and standard deviation from our previous studies and/or pilot studies using 4–5 animals per group. All animal procedures were performed in accordance with protocols approved by the Institutional Animal Care and Usage Committee of New York University School of Medicine.

### Antibodies, intracellular staining and flow cytometry

The following monoclonal antibodies were purchased from eBiosciences, BD Pharmingen or BioLegend: CD3 (145-2C11), CD4 (RM4-5), CD25 (PC61), CD44 (IM7), CD45.1 (A20), CD45.2 (104), CD90.1 (HIS51), CD90.2 (53-2.1), CD19 (1D3), CD45R (RA3-6B2), CD127 (A7R34), CD51 (RMV-7), IA/IE (56-5321-82), CCR6 (3D6), Ncr1 (29A1.4), NK1.1 (PK136), CD62L (MEL-14), CXCR5 (L138D7), TCRβ (H57-597), TCR Vβ6 (RR4-7), Bcl-6 (K112-91), Foxp3 (FJK-16s), RORγt (B2D or Q31-378), T-bet (eBio4B10), IL-17A (eBio17B7) and IFN-γ (XM61.2), CD11c (N418), CD11b (M1/70), CX3CR1 (SA011F11), Ly6c (HK1.4), SIRPa (P84), Ly6G (1A8), CD273 (TY25), Clec12a (5D3), CD103 (M290), XCR1 (ZET), F4/80 (BM8), CCR7 (4B12), CXCR6 (SA051D1), CD40 (HM40-3), 4′,6-diamidino-2-phenylindole (DAPI) or Live/dead fixable blue (ThermoFisher) was used to exclude dead cells.

For transcription factor staining, cells were stained for surface markers, followed by fixation and permeabilization before nuclear factor staining according to the manufacturer’s protocol (Foxp3 staining buffer set from eBioscience). For cytokine analysis, cells were incubated for 5 h in RPMI with 10% FBS, phorbol 12-myristate 13-acetate (PMA) (50 ng/ml; Sigma), ionomycin (500 ng/ml; Sigma) and GolgiStop (BD). Cells were stained for surface markers before fixation and permeabilization, and then subjected to intracellular cytokine staining according to the manufacturer’s protocol (Cytofix/Cytoperm buffer set from BD Biosciences). Flow cytometric analysis was performed on an LSR II (BD Biosciences) or Cytek Aurora (Cytek) or an Aria II (BD Biosciences) and analyzed using FlowJo software (Tree Star).

### Flow cytometry gating strategy

**Hh7-2 gating**: FSC, SSC; Live Dead^-^,singlets, Dump^-^ (B220,TCRgd, Ly6G), MHCII^-^, CD4^+^, TCR*β*^+^, VB6^+^, CD45.1^+^ or CD90.1^+^; **cDC gating**: FSC, SSC; Live Dead^-^,singlets, Dump^-^ (B220, TCR*β*, TCRgd, Ly6G), CD11c^+^ and CD11b^+^, SIRPa^low-moderate^ (remove CD11c^-^, SIRPA^high^); **cDC2 gating** (unless mentioned otherwise): FSC, SSC; Live Dead^-^,singlets, Dump^-^ (B220,TCRgd, Ly6G), CD11c^+^ and CD11b^+^, SIRPa^low-moderate^ (remove CD11c^-^, SIRPA^high^), Clec12a^-^ SIRPa^+^; **migratory cDC2 gating**: FSC, SSC; Live Dead^-^,singlets, Dump^-^ (B220,TCRgd, Ly6G), CD11c^+^ and CD11b^+^, SIRPa^low-moderate^ (remove CD11c^-^, SIRPA^high^), Clec12a^-^ SIRPa^+^, PDL2^+^; **NCR1^+^ ILC3 gating**: FSC, SSC; Live Dead^-^,singlets, Dump^-^ (B220,TCRgd, Ly6G), TCR*β*^-^, CD90+, Il7R^+^, *RORγt* ^+^, CCR6^-^, NCR1^+^; **JC** (using internal staining): FSC, SSC; Live Dead^-^,singlets, Dump^-^ (B220,TCRgd, Ly6G), TCR*β*^-^, ROR*γ*t ^+^, MHCII+ CXCR6^-^, Il7R^-^; **JC** (excluding internal staining): FSC, SSC; Live Dead^-^,singlets, Dump^-^ (B220,TCRgd, Ly6G), TCR*β*^-^, CD11c^low-negative^, MHCII^+^, CCR6^+^,CXCR6^-^, Il7R^-^ ; **ILC3**: FSC, SSC; Live Dead^-^,singlets, Dump^-^ (B220,TCRgd, Ly6G),TCR*β*^-^, CXCR6^+^, Il7R^+^, ROR*γ*t ^+^; **LTi-Like ILC3** (using internal staining): FSC, SSC; Live Dead^-^,singlets, Dump^-^ (B220,TCRgd, Ly6G), TCR*β*^-^, MHCII+, CXCR6^+^, Il7R^+^, ROR*γ*t ^+^, CCR6^+^, CD25^+^; **LTi-Like ILC3** (excluding internal staining): FSC, SSC; Live Dead^-^,singlets, Dump^-^ (B220,TCRgd, Ly6G), TCR*β*^-^, CD11c^low-negative^, MHCII^+^, CCR6^+^,CXCR6^+^, Il7R^+^ Isolation of lymphocytes and APCs

After removal of caecal patches, large intestine tissues were sequentially treated with PBS containing 1 mM DTT at room temperature for 10 min, twice with 5 mM EDTA at 37 °C for 10 min to remove epithelial cells, and then minced and dissociated in digestion buffer (RPMI containing collagenase (1 mg ml−1 collagenase D; Roche), DNase I (100 μg ml−1; Sigma), dispase (0.1 U ml−1; Worthington) and 10% FBS) with constant stirring at 37 °C 55 min. Leukocytes were collected at the interface of a 40%/80% Percoll gradient (GE Healthcare). Lymph nodes were mechanically disrupted for lymphocyte isolation. For isolation of myeloid cells and ILC, lymph nodes were mechanically disrupted with digestion buffer with constant stirring at 37 °C 30 min.

### *H. hepaticus* culture and oral infection

*H. hepaticus* was kindly provided by Dr. James Fox (MIT). Hh was cultured and administrated as was previously described^3^. Frozen stock aliquots of *H. hepaticus* were stored in Brucella broth with 20% glycerol and frozen at −80°C. The bacteria were grown on blood agar plates (TSA with 5% sheep blood, Thermo Fisher). Inoculated plates were placed into a hypoxia chamber (Billups- Rothenberg), and anaerobic gas mixture consisting of 80% nitrogen, 10% hydrogen, and 10% carbon dioxide (Airgas) was added to create a micro-aerobic atmosphere, in which the oxygen concentration was 3∼5%. The micro-aerobic jars containing bacterial plates were left at 37°C for 4 days before animal inoculation. For oral infection, *H. hepaticus* was resuspended in Brucella broth by application of a pre-moistened sterile cotton swab applicator tip to the colony surface.0.3 mL bacterial suspension was administered to each mouse by oral gavage. Mice were inoculated for a second dose after 3 days.

### Adoptive transfer of Hh7-2 TCR transgenic cells

Adoptive transfer of Hh7-2 was done as was previously described^3^, with minor modifications. Recipient mice were colonized with *H. hepaticus* by oral gavage seven days before adoptive transfer. Spleens and lymph nodes from donor Hh7-2 TCRtg mice were collected and mechanically disassociated. Red blood cells were lysed using ACK lysis buffer (Lonza). Naive Hh7-2 T cells were sorted as CD4^+^TCR*β*^+^CD44^lo^CD62L^hi^CD25^−^Vβ6^+^ (HH7-2tg), on the Aria II (BD Biosciences). For analysis of early differentiation, cells were additionally labeled with CFSE (ThermoFisher). Cells were resuspended in PBS on ice and 100K were transferred into congenic isotype-labelled recipient mice by retro-orbital injection. Cells from MLN were analyzed 3 days after transfer and cells from colon LP were analyzed 10-14 days after transfer.

### CITE-seq

CITE-seq and cell hashing were performed as described^44, 45^ with minor modifications. Single-cell suspensions were obtained from digests of C1 MLN of t*dTomato-ON^CD11c^* or *tdTomato-ON^RORɣt^ ; zbtb46-eGFP* mice that had been colonized with *Helicobacter* for 7 days. Cells were sorted on a BD FACSAriaII using a 100-µm nozzle. Dead cells as well as T cells and B cells were gated out using DAPI, TCR*β*, TCR*γδ* and B220 antibodies. From t*dTomato-ON^CD11c^* mice, the tdTomato+ population was collected separately from two mice. From *tdTomato-ON^RORɣt^ ; zbtb46-eGFP* mice, we collected 3 populations from two separate mice: GFP^+^, GFP^+^ tdTomato^+^ and tdTomato^+^. Sorted cells were stained separately with hashing antibodies (Biolegend)^45^. After removal of excess hashing antibodies, we combined the samples and stained them with CITE-seq antibodies, conjugated using iEDDA click chemistry to barcode oligos as described before^46^. In addition, we included some commercially available Totalseq-A antibodies for CD11c, CD90.2, CD185, CD51 and CD127 (Biolegend). Post-sorting and staining, cells were run through the standard 10x Chromium (v3) protocol up until cDNA amplification, with the following modification: For the cDNA PCR step, 0.2uM of ADT additive primer (5’CCTTGGCACCCGAGAATTCC) and 0.2uM of HTO additive primer (5’GTGACTGGAGTTCAGACGTGTGCTC) were added to the cDNA amplification master mix. Post cDNA amplification, a 0.6X SPRI cleanup was performed to separate the cDNA fraction (on beads) from the smaller ADT and HTO fractions (in supernatant). The cDNA fraction was converted into a 3’ tag gene expression library according to the 10x Genomics Single Cell Genomics Protocol (v3). Supernatant from cleanup was kept for ADT and HTO preparation.

To the supernatant, another 1.4X SPRI was added to bring the total SPRI concentration to 2X. After washing the beads in 80% ethanol and eluting in water, a second round of 2X SPRI cleanup was performed to remove any residual primer carryover from the cDNA PCR. Post cleanup, eluate was taken into ADT PCR amplification (using TruSeq Small RNA RPIx primer (5’CAAGCAGAAGACGGCATACGAGXXXXXXXXGTGACTGGAGTTCCTTGGCACCCGAGAAT TCCA) and SI PCR primer (5’AATGATACGGCGACCACCGAGATCTACACTCTTTCCCTACACGACGCTC)), and into hashtag amplification (using TruSeq D7xx (5’CAAGCAGAAGACGGCATACGAGATXXXXXXXXGTGACTGGAGTTCAGACGTGTGC) and SI PCR primers). Libraries were pooled and sequenced on a 100 cycle Novaseq S1 flowcell, with the configuration of 30 base pairs for R1, and 92 base pairs for R2. Additional protocol details can be found for CITE-seq and cell hashing at www.cite-seq.com.

Post sequencing, gene expression count matrices were generated using cellranger version 5.0 using the refdata-gex-mm10-2020-A reference library provided by 10x Genomics, with the additional sequences of cre, eGFP, and tdTomato. Counts matrices for hashtags and antibodies were generated using CITE-seq-Count version 1.4.4. Downstream analysis was then performed in R using Seurat.

Quality control and doublet removal: We initially selected all cells that were detected in our RNA- seq, cell hashing, and ADT libraries. We removed cells with < 700 detected genes, but also removed cells which had an aberrantly high number of genes (more than 5,000 genes) and a high percentage of mitochondria genes (more than 6%). Additionally, we removed the cells which were attached by the clumps of antibodies and had a too high number of ADT or HTO UMIs (more than 5,000 ADT UMI and 4,000 HTO UMI). We used our previously described hashing-based doublet detection strategy^45^, implemented in HTODemux, to identify doublets that represent two or more cells representing different samples. Only Singlet cells were used for the downstream analysis.

Multimodal analysis: We normalized RNA data using SCTransform^47^ and applied the centered- log ratio (CLR) transformation to normalize ADT data within each cell. We used principal component analysis (PCA) to reduce dimensionality of both datasets. Then we took both top 20 RNA and protein PCA dimensions as the input of weighted nearest neighbors (WNN) method^48^ to construct the multimodal weighted KNN graph. To cluster our multimodal dataset, we first used the weighted KNN graph to generate a shared nearest neighbor graph (SNN) and then apply the graph-based smart local moving (SLM) algorithm (https://doi.org/10.1140/epjb/e2013-40829-0) on this SNN graph to find clusters with 0.8 resolution. We performed differential expression on all pairs of clusters for both RNA and protein markers, and merged clusters that did not exhibit clear evidence of separation. All samples were clustered together and separated later for further analysis as indicated.

### CITE-seq data projection in Flowjo

For gating analysis, scaled and normalized ADT counts together with cluster identity, UMAP and UWnn coordinates were exported into a csv format that was uploaded in FlowJo.

### Single cell RNAseq of Aire^+^ cells

Four week-old Adig mouse lymph nodes (cervical, brachial, axillary, inguinal, and mesenteric) (n=3) were pooled in digestion medium consisting of RPMI 1640 with 2% fetal bovine serum (FBS) (Sigma-Aldrich), deoxyribonuclease (DNase) (100 µg/ml; Roche) and Liberase (50 µg/ml; Roche), minced and agitated at 37°C for 30 min, and passed through a 70-µm filter. Cells were resuspended for magnetic column enrichment (Miltenyi LD column depletion with streptavidin microbeads and biotinylated antibodies against B220, Ter119, TCRbeta, CD3e). Cells were then either processed directly for 10x single-cell analysis as reference sample or sorted by flow cytometry for all live GFP+ cells. Cells were sorted into PBS with 0.04% bovine serum albumin (BSA). Cell viability and counts were evaluated with Vi-CELL XR (Beckman Coulter), and samples with viability >85% were used for sequencing.

For analysis, sequencing files were aligned to the mm10 mouse reference genome with the 10x Genomics Cell Ranger (v3.1.0) count method using the default parameters. Raw count files were then processed in Python (v3.9.7), removing doublets from each sample using Scrublet^49^ with a doublet score threshold of 0.3. Samples were merged and filtered, removing genes with no counts, and retaining cells with 600 to 5,000 genes and 1,000 to 30,000 counts, leaving a total of 35,797 cells and 22,979 genes in the dataset. The raw counts were used to train a model of gene expression using scvi-tools (v0.14.6) with each sample as a batch key. This model was used to generate normalized expression values for all genes scaled to a library size of 100,000 and create a tSNE representation of the data using Scanpy^50^ (v1.8.2) with the default parameters. Leiden clustering with a resolution of 0.8 gave 29 clusters which were assigned cell types after removing three low quality clusters. For targeted analysis of JC, the data was subset on the three JC clusters and a UMAP was created using Scanpy with a minimum distance of 0.5 and spread of 1. Differential gene expression of JC clusters was performed using scvi-tools^51^, filtering on genes with a bayes factor greater than 3, mean log fold change greater than 0, and proportion of cells with non-zero expression greater than 0.1. The z-score of the average expression of the top 10 DE genes from each cluster was used for visualization. For low-dimensional embeddings of feature plots, scVI normalized expression was used, clipping the top and bottom 1% of expression values in the full dataset to the maximum and minimum of the color scale to prevent outliers from skewing the visualization. To display expression of key features in dotplots, log-normalized and scaled counts for each gene were averaged and standardized across cell types to have a value ranging from 0 to 1.

### RNAscope

Fresh-frozen spleen and lymph nodes were sectioned at 8um and then fixed overnight at 4°C in 10% neutral buffered formalin (Thermo Fisher Scientific, Waltham, MA) before proceeding with an RNAscope RED 2.5 HD Chromogenic Assay kit (Advanced Cell Diagnostics, Newark, CA) for detection of Aire mRNA. DapB probe was used as negative control and Polr2a probe was used as positive control. Semi-quantitative scores were determined in a blinded fashion based on the number of Aire^+^ cells per section.

### Integrin *β*8 (ADWA-11) blocking Ab

We injected IP 200ug of ADWA-11 into mice colonized with Hh for 7 days. On the same day we adoptively transferred 100K CFSE-labeled naive Hh7-2 cells and tested their proliferation and differentiation in the C1 MLN, 3 days after the transfer.

### Generation of bone marrow (BM) chimeric reconstituted mice

To generate chimeric mice, 4-5 week old CD45.1 mice were irradiated twice with 500 rads/mouse at an interval of 2-5 h (X-RAD 320 X-Ray Irradiator). A day after, bone marrow (BM) mononuclear cells were isolated from donor mice, as indicated in each experiment, by flushing the femur bones. Red blood cells were lysed with ACK Lysing Buffer, and lymphocytes were depleted for Thy1.2 using magnetic microbeads (Miltenyi). BM cells were resuspended in PBS and a total 3-4 x 10^6^ BM cells were injected intravenously into the irradiated mice. In case of mixed BMC reconstitution, a ratio of 1:1 was used. Mice were kept for a week on broad spectrum antibiotics (1 mg/mL sulfamethoxazole and 0.2 mg/ mL trimethoprim), followed by microbiome reconstitution by fecal gavage. Mice were reconstituted for 1-2 months before Hh colonization. After 7 weeks, peripheral blood samples were collected and analyzed by FACS 7 to check for reconstitution.

### Intravital multiphoton microscopy

Naive Hh7-2 T cells were isolated from *Nur77-eGFP* Hh7-2tg mice, labeled with Cell tracker dye (eBioscience Cell Proliferation Dye eFluor 450), and transferred into *mKate2-ON^zbtb46^;RORγt- eGFP* mice that had been colonized with Hh for 6 days. Fifteen hours following adoptive transfer of Hh T cells, mice were euthanized and C1 MLN were immediately isolated and mounted in cold RPMI with 10% FCS for intravital multiphoton microscopy. Image stacks were acquired with an Olympus multiphoton FVMPE-RS system equipped with both InSight X3 and Mai Tai Deepsee (Spectra-Physics) tunable Ti:Sapphire lasers. To acquire serial optical sections, a laser beam (780 nm for eFluor™ 450 and 940 nm for simultaneous excitation of eGFP and mKate2) was focused through a water immersion lens (N.A. 1.05; Olympus) and scanned with a field of view of 0.5 mm^2^, at 600 Hz. Z-stacks were acquired in 2 mm steps to image a total depth of 150-200 mm of tissue.

### Image analysis

Raw image stacks were imported into Fiji (NIH) for T cell colocalization analysis. Provided images are presented as a maximal projection of 3–6 mm optical sections. For visualizing individual labelled cells expressing both the mKate2 and eGFP, the brightness and contrast were adjusted accordingly to single positive green (eGFP) and red (mKate2) cells. Adoptively transferred Hh7- 2tg T cells were identified via positive labeling with cell proliferation dye eFluor 450. Primed Hh7- 2tg T cells were identified via expression of the *Nur77*-*eGFP* reporter. Cell identity was scored by a combination of both fluorescent reporter expression as well as cell morphology. Specifically, cells expressing mKate2 with a dendritic cell shape were scored as cDC, while cells expressing both *RORɣt*-*eGFP* and mKate2 (or *eGFP* alone) with an amoeboid (non-spherical) cell shape were scored as ILC3. T cell interactions with cDC or ILC3 was strictly measured as direct (<1 micron) colocalization of cells with respective fluorescent and cell morphology combinations.

### Statistical analysis

For animal studies, mutant and control groups did not always have similar standard deviations and therefore an unpaired two-sided Welch’s t-test was used. Error bars represent ± s.d. Animal sample size estimates were determined using power analysis (power = 90% and α = 0.05) based on the mean and s.d. from our previous studies and/or pilot studies using 4–5 mice. No samples were excluded from analysis. For analysis of ILC3 counts in MLN and LP of chimeric mice, a paired two-sided t-test was used.

### Data availability

Data generated for this project are available at the Gene Expression Omnibus with the accession code GSE190372 and XXX. Published data GSE176282 was used for analysis.

### Code availability

All code used for analysis in this manuscript is available at https://github.com/nygctech/Kedmi-CITEseq.

---

**Extended Data Fig. 1.**
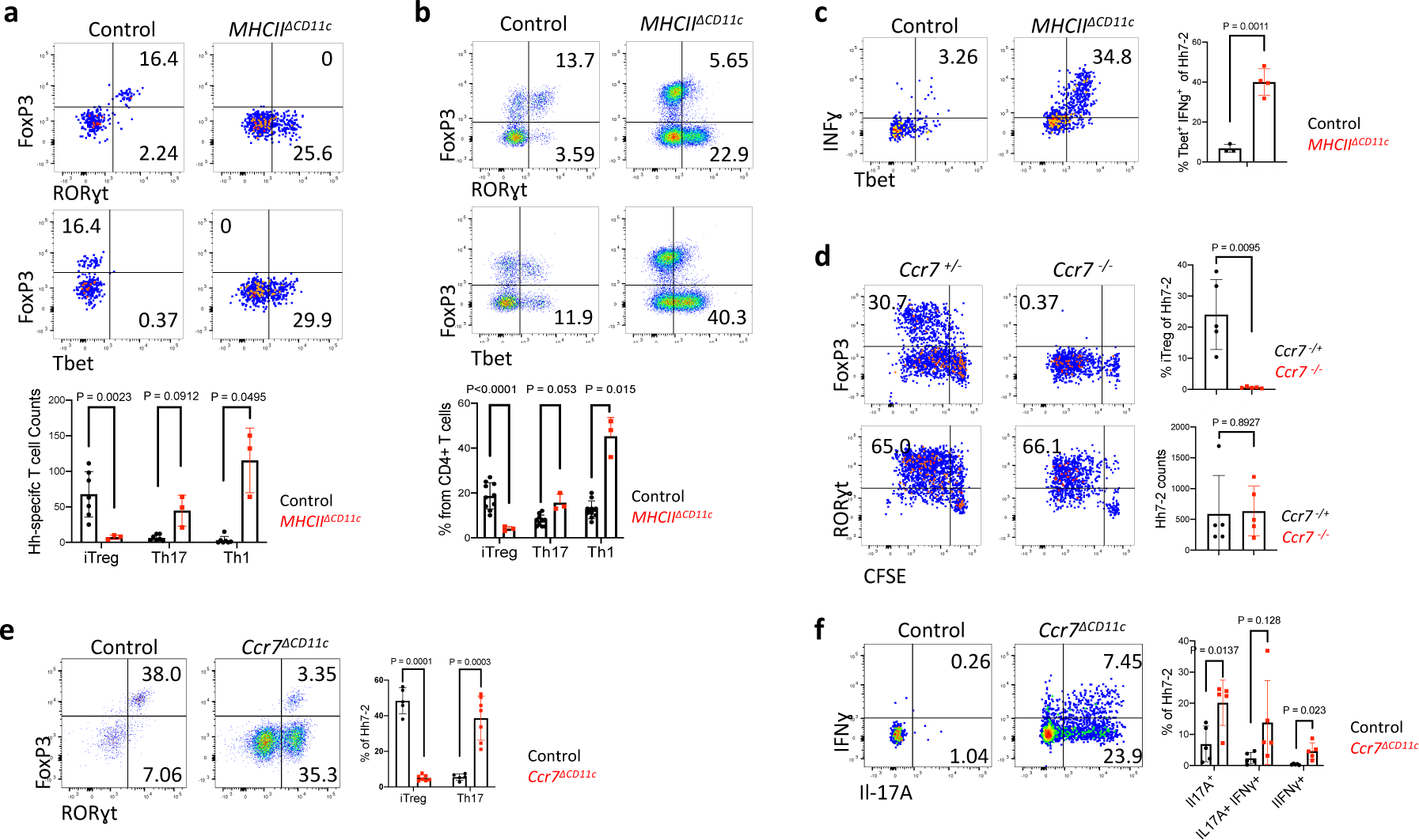
Cells targeted by CD11c-Cre and consequences for Hh-specific T cell differentiation. **a,** Phenotype of Hh7-2 TCR transgenic T cells in the colon lamina propria at 10 days after transfer into Hh-colonized *MHCII^ΔCD11c^* (n=3) and control mice (n=7), as indicated. **b,** Phenotype of host CD4^+^ T cells from mice in (a); *MHCII^ΔCD11c^* (n=3) and control mice (n=10), as indicated. **c,** Cytokine profile of Hh7-2 T cells shown in (a); *MHCII^ΔCD11c^* (n=4) and control mice (n=3). **d,** Proliferation and differentiation of Hh-specific iTreg and Th17 cells in the MLN of *Ccr7^-/-^* (n=5) and littermate control mice (n=5). CFSE-labeled Hh7-2 T cells were analyzed at 3 days following their adoptive transfer into Hh-colonized mice. Data summarize two independent experiments. **e-f,** Transcription factor (e) and intracellular cytokine (f) profiles of Hh7-2 T cells in the large intestine of *Ccr7^ΔCD11c^* (n=7 or 5, for transcription factors and cytokines, respectively) and littermate control (n=5) mice, at 10 days after adoptive transfer. Data summarize two independent experiments. Representative flow panels and aggregate data are shown for each analysis. All statistics were calculated by unpaired two-sided Welch’s t-test. Error bars denote mean ± s.d. *p*-values are indicated in the figure.

**Extended Data Fig. 2.**
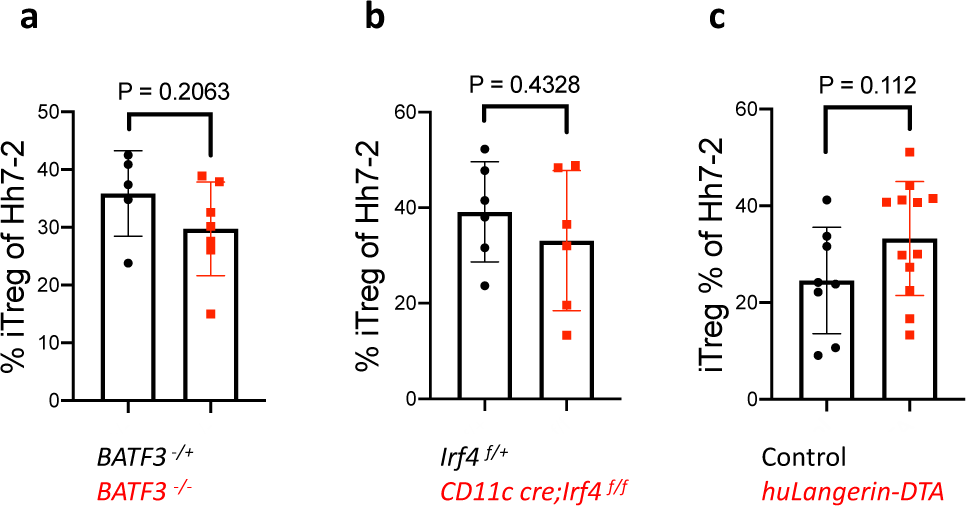
cDC1 or cDC2 are not required for iTreg differentiation. **a-c,** Proportion in MLN of Hh7-2 with the iTreg phenotype at 3 days after transfer into *BATF3^-/-^* (a) (n=7), *IRF4^ΔCD11c^* (b) (n=6), and *huLangerin (CD207)-DTA* (c) (n=12) mice (red) and indicated littermate controls (black). Data summarize at least two independent experiments. All statistics were calculated by unpaired two-sided Welch’s t-test. Error bars denote mean ± s.d. *p*-values are indicated in the figure.

**Extended Data Fig. 3.**
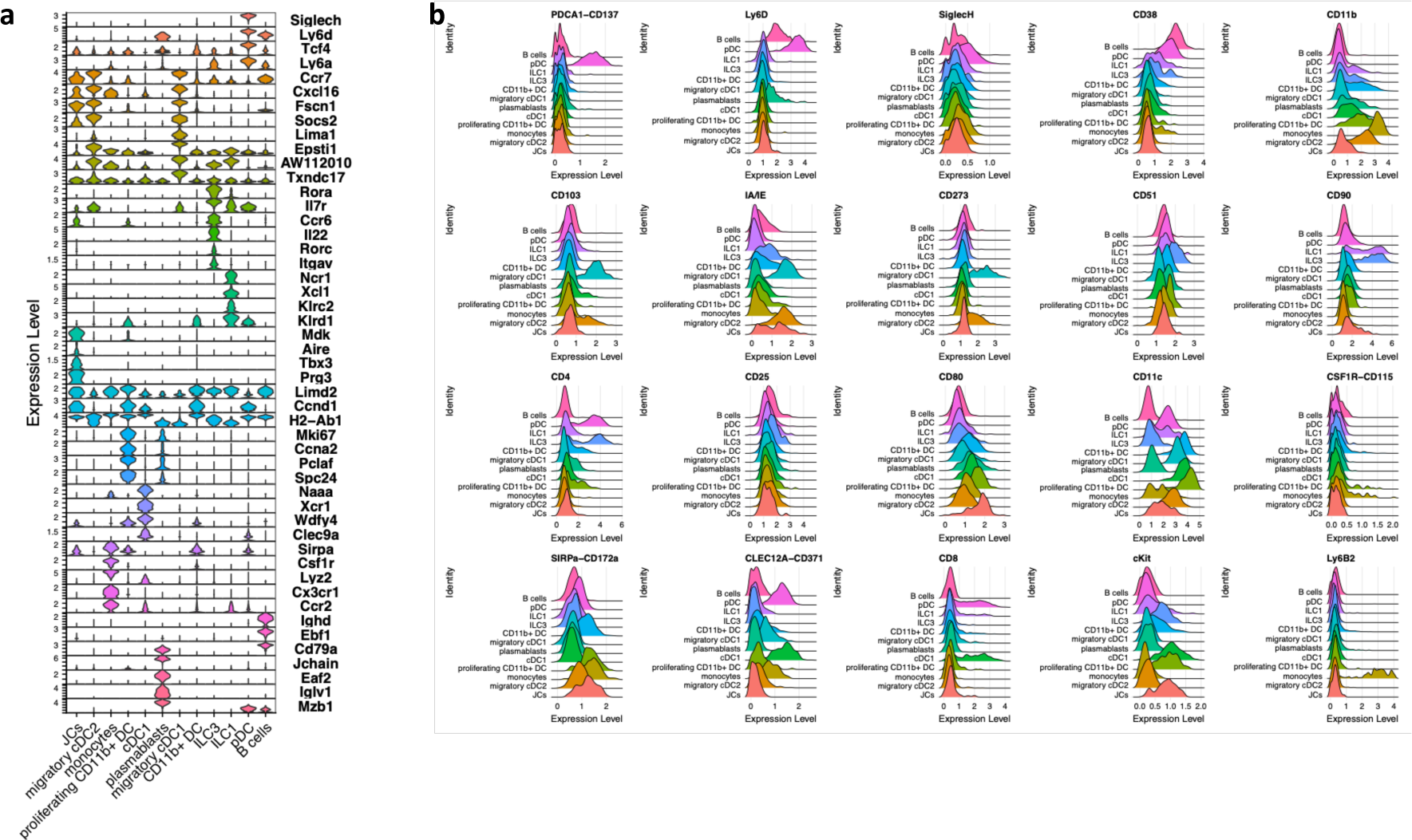
Identification of CITE-seq-assigned clusters of sorted *tdTomato- ON*^ΔCD11c^ fate-mapped cells. **a,** Stacked violin plots for selected (curated) and top DEG (data- driven) of tdTomato^+^ cells sorted from MLN of Hh-colonized mice. **b,** Ridge Plot of cell surface markers for each cluster.

**Extended Data Fig. 4.**
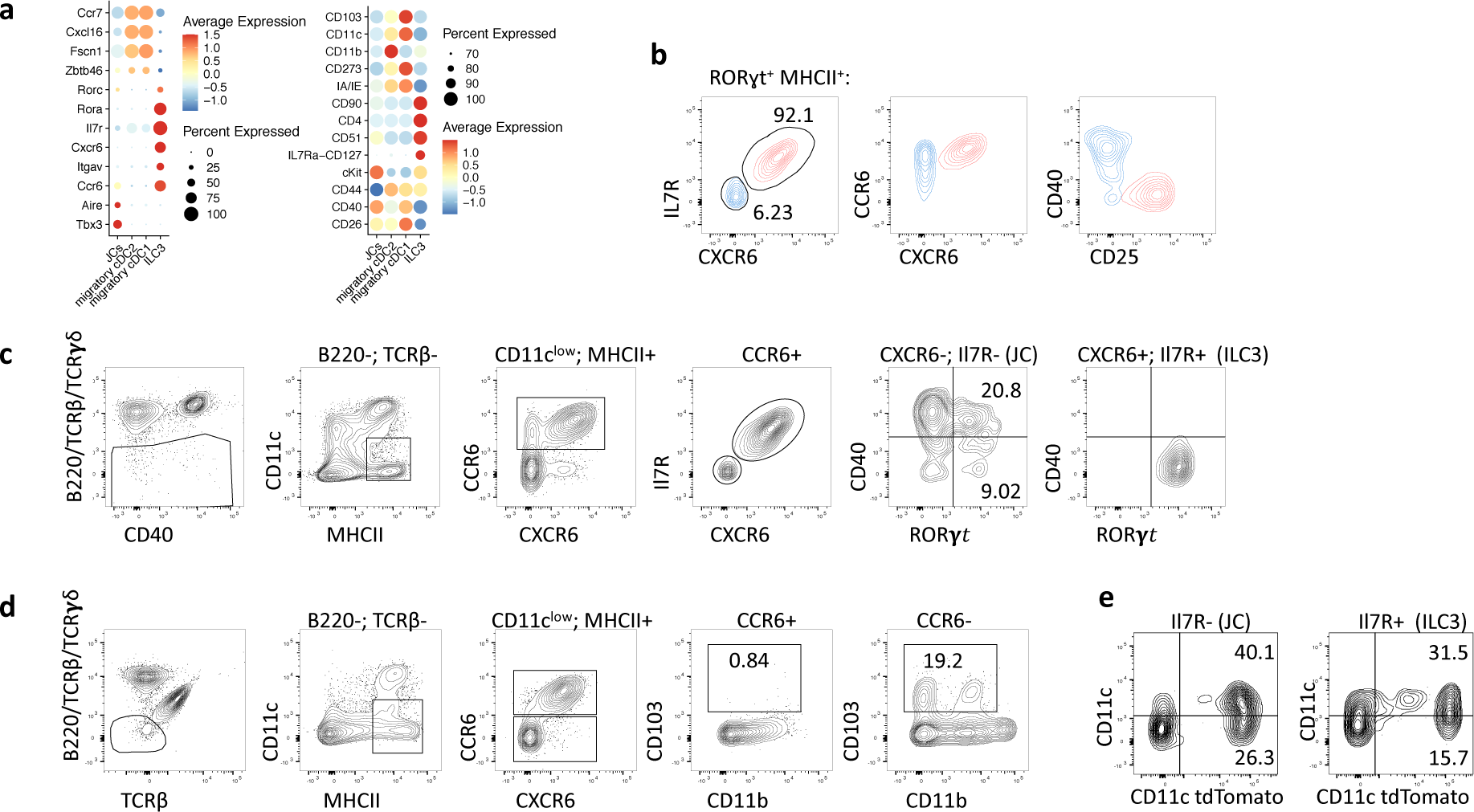
Phenotypic discrimination of ILC3 and JC. **a,** Dot plots for selected (curated) DEG and cell surface markers for the indicated clusters, obtained from CITE-seq analysis of *tdTomato-ON^ΔCD11c^* fate-mapped cells. **b,** Flow cytometry profiling of CXCR6, CD127(IL-7R), CCR6 CD25 and CD40 on ILC3 (red) and JC (blue), pre-gated on TCR*β*^-^, TCR*γδ*^-^, B220^-^, ROR*γ*t^+^, MHCII^+^. **c,** gating strategy for JC using cell surface staining as indicated. **d,** Flow cytometry profiling of JC and DC markers, showing that migratory cDC are excluded from CD11c^low^ CCR6^+^ gating. **e,** TdTomato levels in ILC3 (TCR*β*^-^, TCR*γδ*^-^, B220^-^, MHCII^+^ CCR6^+^, Il7R^+^) and JC (TCR*β*^-^, TCR*γδ*^-^, B220^-^, MHCII^+^ CCR6^+^, Il7R^-^) from the MLN of Hh-colonized *tdTomato-ON^CD11c^* fate-map mice.

**Extended Data Fig. 5.**
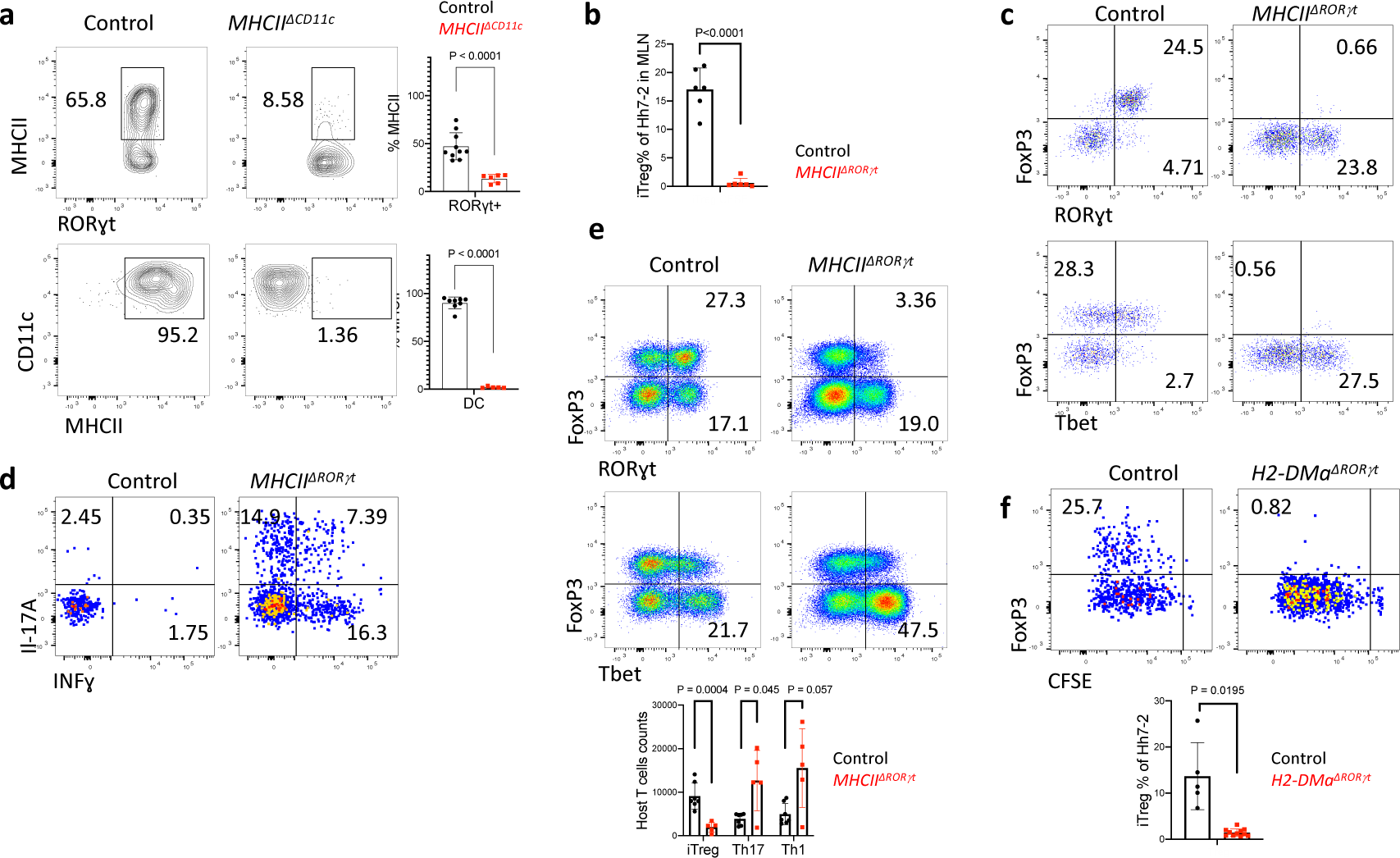
Antigen presentation by ROR*γ*t^+^ cells is required for microbiota-induced iTreg cell differentiation. **a,** MHCII expression in ROR*γ*t^+^ cells (top) and DC (bottom) from the MLN of Hh-colonized *MHCII^ΔCD11c^* mice (n=6 and 5) and littermate controls (n =10 and 8). ROR*γ*t^+^ cells were gated as TCR*β*^-^, TCR*γδ*^-^, B220^-^, ROR*γ*t^+^; DC were gated as TCR*β*^-^, TCR*γδ*^-^, B220^-^, CD90^-^, CD11c^high^. b, Bar graph showing frequency of iTreg among Hh7-2 T cells, measured as in Figure 2e. c-d, Representative dot plots showing Hh7-2 T cell differentiation (c) and cytokine (d) profiles in colon lamina propria at 22 days after adoptive transfer into Hh-colonized *MHCII^ΔRORγt^* and littermate controls. e, Representative and aggregate data of transcription factor profiles of host CD4^+^ T cells in colon lamina propria of mice shown in (c) and (d). **f,** Hh7-2 cell proliferation and differentiation in the MLN of *H2-DMa^ΔRORγt^* (*RORγt-Cre; H2-Dma^f/f^*) (n=11) and littermate controls (*RORγt-Cre; H2-DMa^+/f^*) (n=5) at 3 days after transfer of CFSE labeled naïve Hh7-2, cell proliferation and FoxP3 were assessed in cells isolated from C1 MLN. Representative flow cytometry (left) and aggregate data from multiple animals (right). Data summarize two independent experiments. All statistics were calculated by unpaired two- sided Welch’s t-test. Error bars denote mean ± s.d. *p*-values are indicated in the figure.

**Extended Data Fig. 6.**
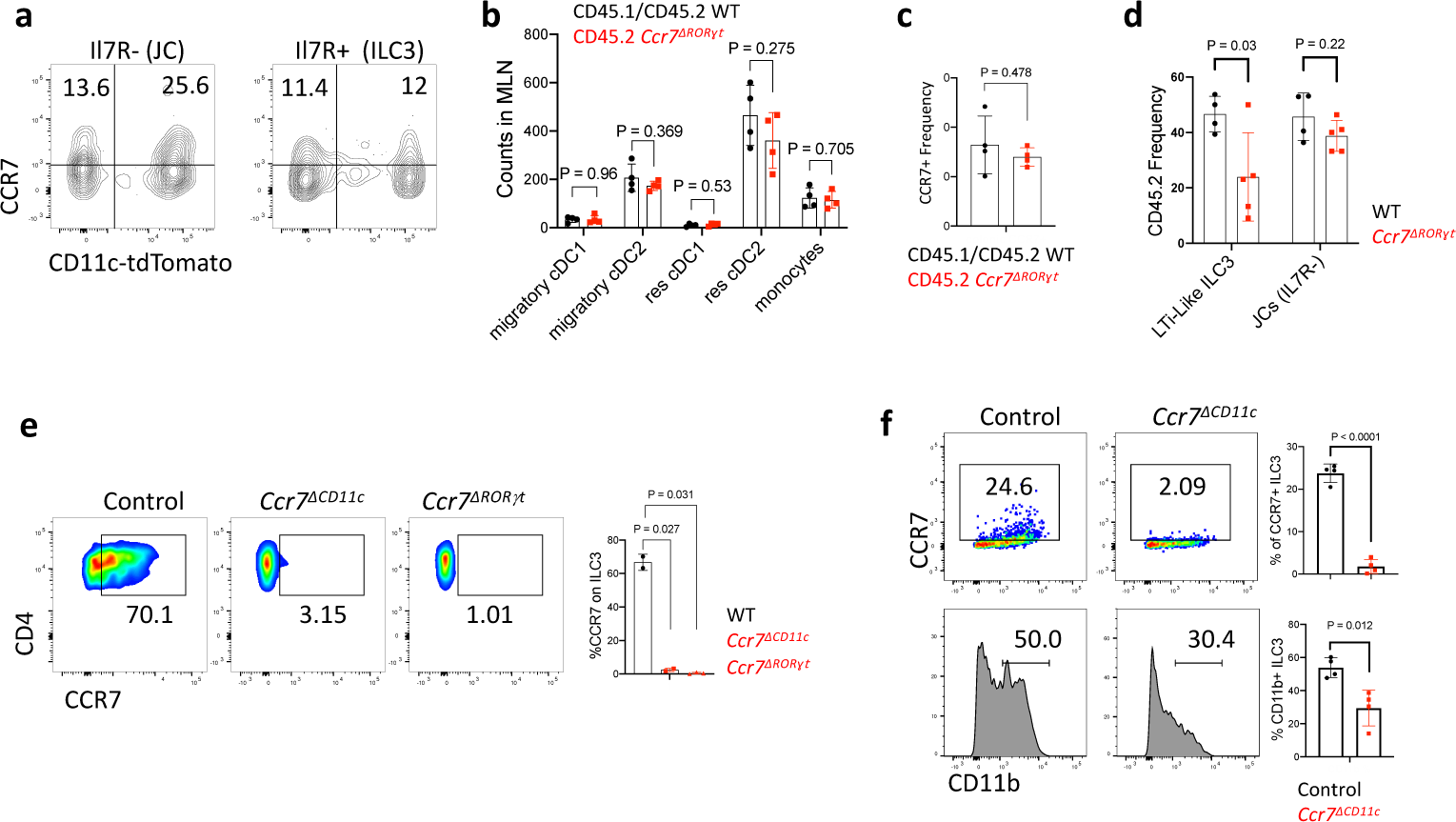
Differential requirements for CCR7 in iTreg and effector Th17 cell differentiation. **a,** Cell surface expression of CCR7 on *CD11c-Cre* fate-mapped ILC3 (TCR*β*^-^, TCR*γδ*^-^, B220^-^, MHCII^+^, CCR6^+^, IL-7R^+^) and JC (TCR*β*^-^, TCR*γδ*^-^, B220^-^, MHCII+, CCR6+, IL-7R^-^) in the MLN. **b-c,** Analysis of DC counts in MLN (b) and large intestine (c) of WT and *Ccr7^ΔRORγt^* mixed bone marrow chimeric mice described in Figure 3c. Counts in the MLN of DC subsets derived from bone marrow (b); frequencies of CCR7^+^ among total colonic DCs (c) (n=4). Statistics were calculated using paired two-sided t-test. **d,** Analysis of CD45.2 frequencies within donor cells is presented for ILC3 (TCR*β*^-^, TCR*γδ*^-^, B220^-^, MHCII^+^, ROR*γ*t ^+^, IL-7R^+^) and JC (TCR*β*^-^, TCR*γδ*^-^, B220^-^, MHCII^+^, ROR*γ*t ^+^, IL-7R^-^) in the MLN of WT and *Ccr7^ΔRORγt^* mixed bone marrow chimeric mice described in Figure 3c. **e,** Cell surface expression of CCR7 in colonic ILC3 (TCR*β*^-^, TCR*γδ*^-^, B220^-^, CD90^+^, ROR*γ*t^+^, CD25^+^, CD4^+^) from *Ccr7^Δ^*^ROR^*^γ^*^t^ (n=3), *Ccr7^ΔCD11c^* (n=2) and control Hh-colonized mice (n=2). **f,** Cell surface expression of CCR7 and CD11b in ILC3-gated MLN cells (TCR*β*^-^, TCR*γδ*^-^, B220^-^, IL-7R^+^, CCR6^+^, CD25^+^) from *Ccr7^ΔCD11c^* (n=4) and control Hh-colonized mice (n=4).

**Extended Data Fig. 7.**
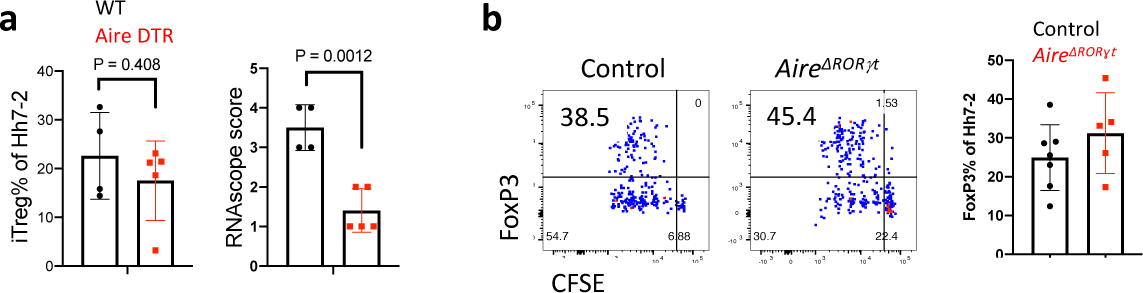
Analysis of Aire^+^ JC function in differentiation of Hh-specific iTreg cells. **a,** Lethally irradiated mice were reconstituted with BM cells from CD45.2 *Aire-DTR* or CD45.2 WT mice. One month after reconstitution, mice were colonized with Hh, and one week later were treated with Diphtheria toxin (DT, Sigma-Aldrich) for 3 sequential days (at a dose of 25 ng/g mice). CD45.1/CD45.2 CFSE-labeled Hh7-2 T cells (1 x 10^5^) were transferred intravenously into the mice on the first day of DT treatment. Bar graph of proportion of proliferating Foxp3^+^ Hh7-2 T cells in the MLN of mice reconstituted with Aire-DTR BM (n=5) or with WT BM (n=4) (left); *Aire* mRNA in the spleen of the treated mice was blindly scored using RNAscope analysis. **b,** Proliferation and differentiation of CFSE-labeled Hh7-2 T cells in the MLN of *RORγt-Cre;Aire^f/f^* (n=5) and control *Aire^+/f^* littermates (n=7) at 3 days after adoptive transfer. Data summarize three independent experiments. All statistics, except for b and c, were calculated by unpaired two-sided Welch’s t-test. Error bars denote mean ± s.d. *p*-values are indicated in the figure.

**Extended Data Fig. 8.**
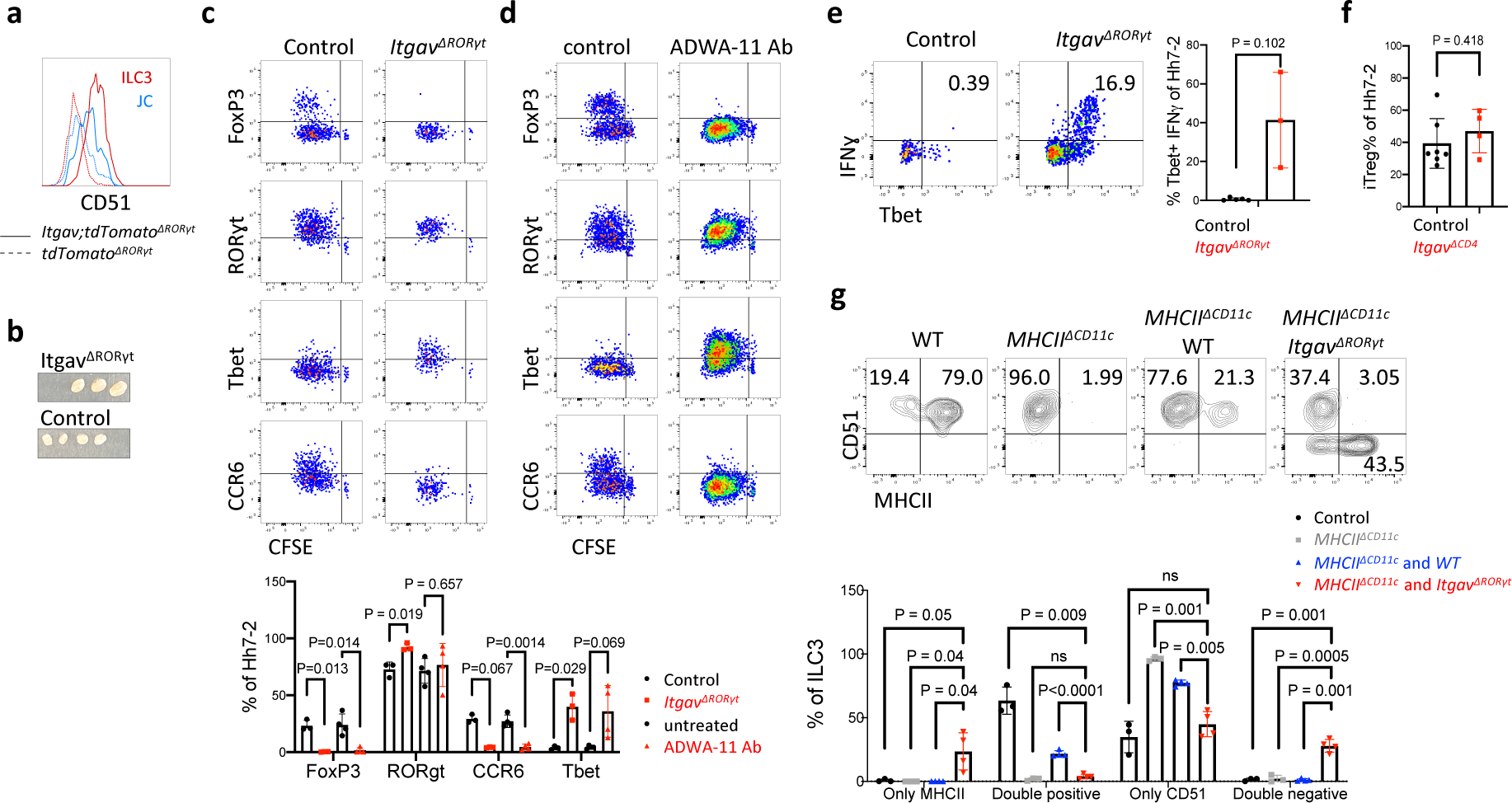
Effect of integrin *α*v*β*8 blockade or *α*v inactivation on microbiota- dependent T cell differentiation. **a,** Expression of integrin *α*v (CD51) in fate-mapped ROR*γ*t^+^ cell subsets from MLN of wild type and *Itgav^ΔRORγt^* mice. **b,** C1 MLN from *Itgav^ΔCD11c^* (n=3) and littermate controls (n=4), 10 days after Hh colonization. **c-d,** Flow cytometry profiling of transcription factors and CCR6 in proliferating CFSE-labeled Hh7-2 in the MLN at 3 days after adoptive transfer into *Itgav^ΔRORγt^* (n=3) and littermate control mice (n=3) (b) or into mice treated with ADWA11(n=4) (as in Fig. 4a) or untreated control littermates (n=4) (c). Summary data of results in (b) and (c) are shown below. **e,** Intracellular IFN*γ* and T-bet expression in PMA/Ionomycin-stimulated Hh7-2 T cells isolated from colon lamina propria of *Itgav^ΔRORγt^*(n=3) and control littermates (n=5), 10 days after adoptive transfer. **f,** Frequency of iTreg cells among proliferating Hh7-2 in the MLN at 3 days after adoptive transfer into *Itgav^ΔCD4^* (n=4) and control littermates (n=7). Data summarize two independent experiments. **g,** Integrin *α*v and MHCII cell surface expression in ILC3 (gated as TCR*β*^-^, TCR*γδ*^-^, B220^-^, ROR*γ*t^+^, CD90^+^, CD25^+^ CD45.2^+^) isolated from MLN of bone marrow chimeric mice, reconstituted with different combinations of donor cells as indicated and colonized with Hh for 10 days. Data summarized below for control (n=3), *MHCII^ΔCD11c^* (n=3), *MHCII^ΔCD11c^* and WT (n=4), and *MHCII^ΔCD11c^* and *Itgav^ΔRORγt^* (n=4) reconstituted mice. All Statistics were calculated by unpaired two-sided Welch’s t-test. Error bars denote mean ± s.d. *p*-values are indicated in the figure.

**Extended Data Fig. 9.**
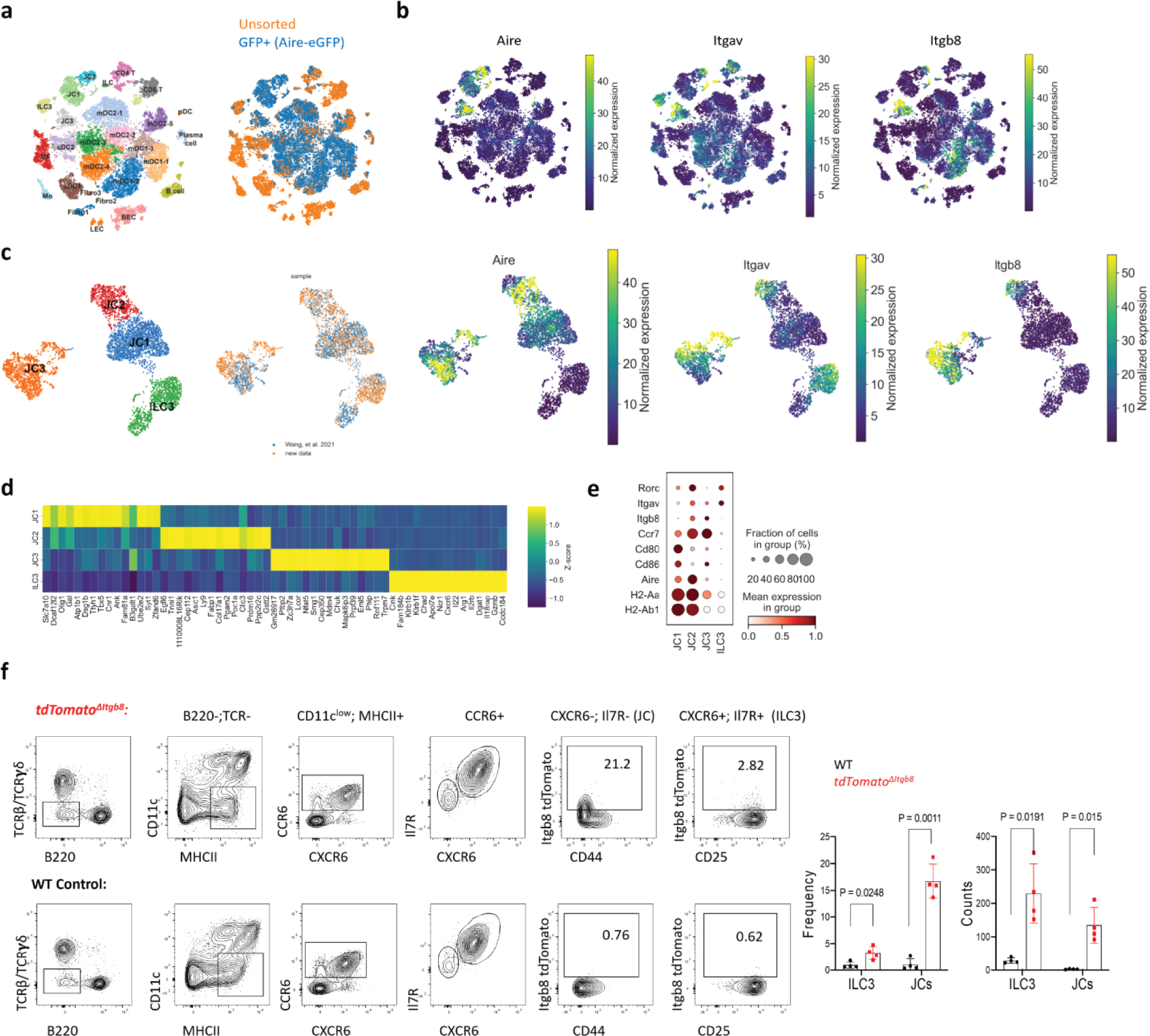
*Itgav and Itgb8* expression in ILC3 and JC. **a**, tSNE plot with Leiden clustering of scRNAseq of pooled GFP^+^ sorted and unsorted cells, as indicated, from pooled lymph nodes of Adig mice^32^. **b,** tSNE feature plots showing *Aire, Itgav, and Itgb8* levels in the cell clusters. **c,** UMAP plot of Aire+ JC and ILC3 populations from pooled datasets as indicated with associated feature plots. **d,** top differentially expressed genes per pseudobulk cluster in (c), shown by heatmap. **e,** dot plot of selected genes in JC and ILC3 clusters. **f,** Representative flow cytometry (left) and aggregate results (right) of tdTomato^+^ JC and ILC3, gated as indicated, in C1 MLN of *tdTomato^ΔItgb8^* mice^33^ (n=4) and littermate controls (n=4). Aggregate data (right) show percent tdTomato+ cells among total ILC3 and JC and number of reporter-positive cells in the C1 MLN of each mouse. All statistics were calculated by unpaired two-sided Welch’s t-test. Error bars denote mean ± s.d. *p*-values are indicated in the figure.

**Extended Data Fig. 10.**
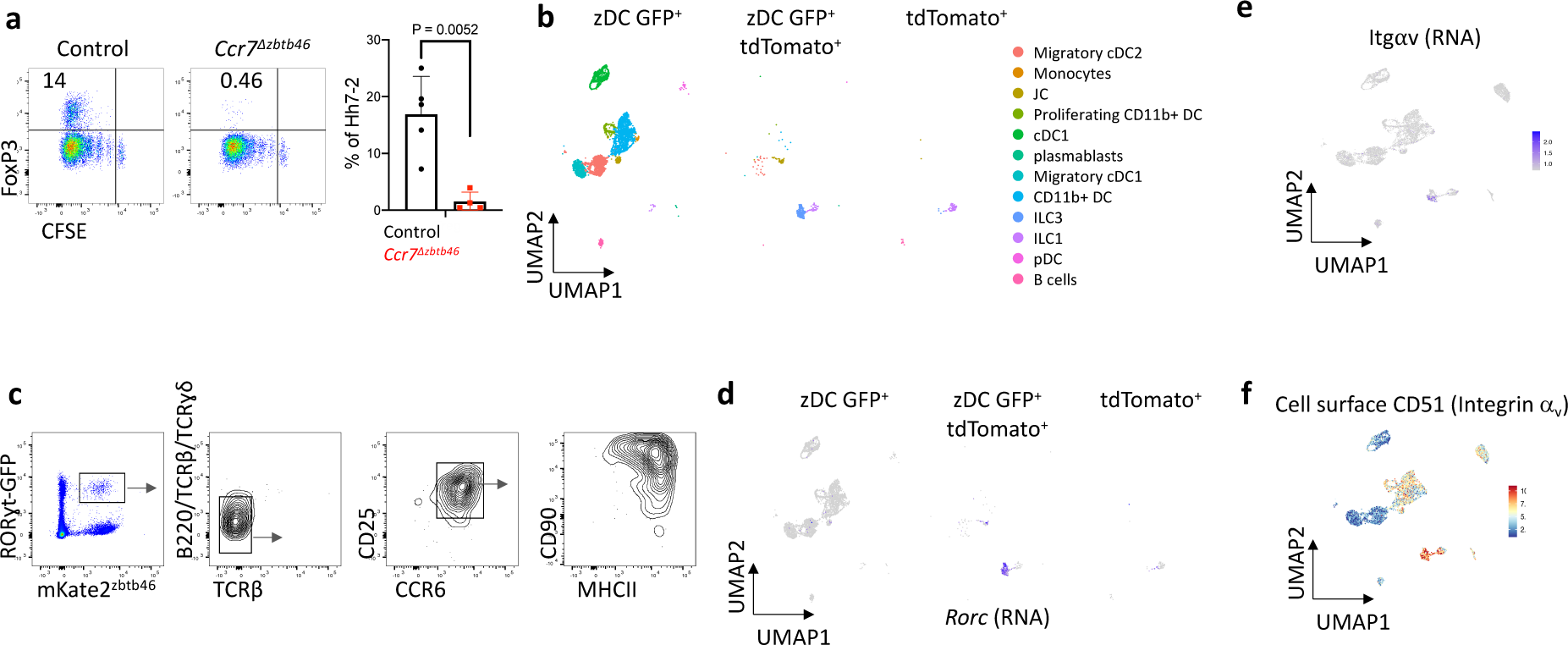
Analysis of ROR*γ*t-expressing cells in the MLN. **a,** Hh7-2 proliferation and differentiation in MLN of *Ccr7^Δzbtb46^* (n= 4) and control littermates (n=5), at 3 days after transfer of the naïve cells. Data in the right panel summarize three independent experiments. **b,** UMAP visualization of CITE-seq datasets obtained from 3 distinct sorted populations (GFP^+^, GFP^+^ tdTomato^+^ and tdTomato^+^) isolated from C1 MLN of *Zbtb46-eGFP ; tdTomato-ON^ΔRORγt^* mice (n=2), analyzed by the WNN method. **c,** Flow cytometry analysis of fate-mapped C1 MLN cells from *RORγt-eGFP;mKate2-ON^Δzbtb46^* mice, gated for the indicated cell subsets. **d-e,** Feature plot showing *Rorc* (d) and integrin *α*v (e) levels in the cell clusters identified in the CITE-seq analysis shown in (b); Positive cells are layered in front. All statistics were calculated by unpaired two-sided Welch’s t-test. Error bars denote mean ± s.d. *p*-values are indicated in the figure.

**Extended Data Fig. 11.**
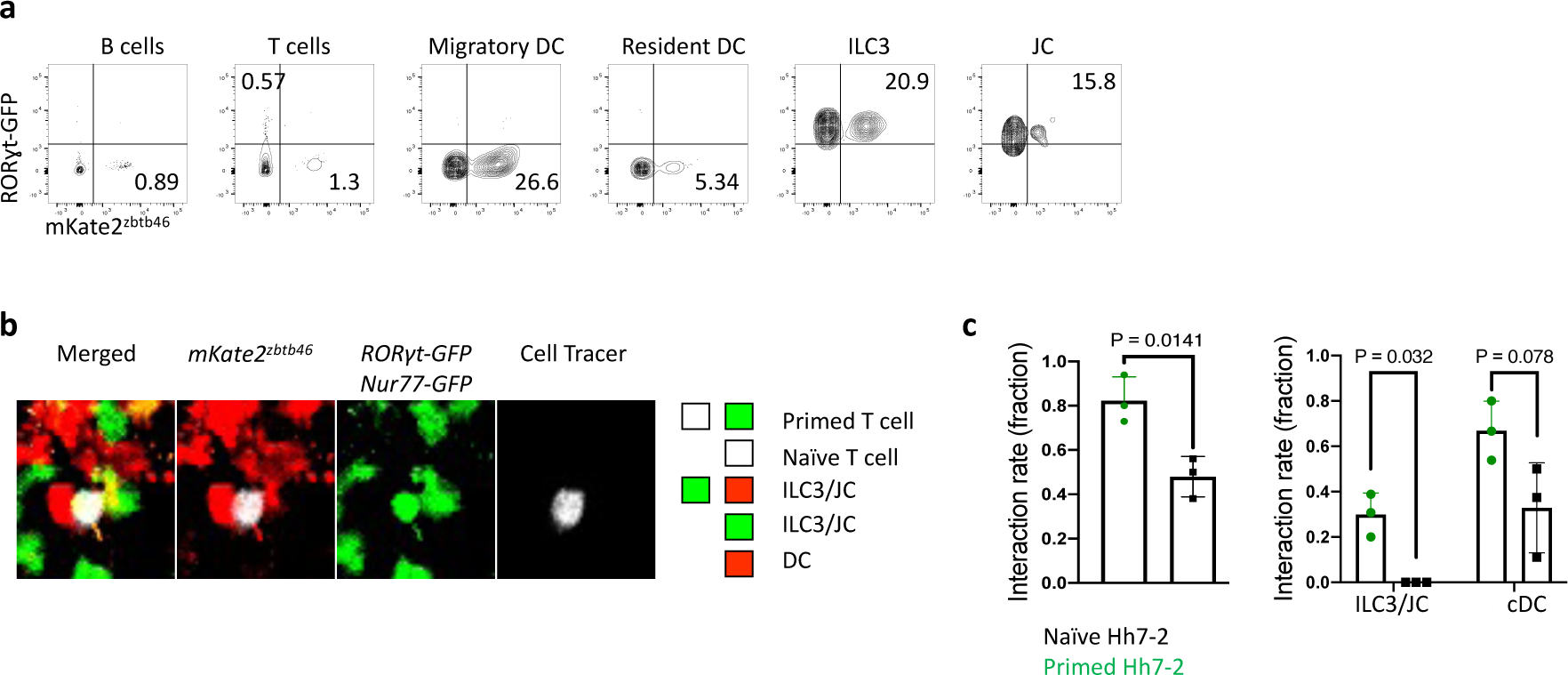
Intravital tracking of ROR*γ*t-expressing cells and DC interactions with Hh-specific T cells during priming in the MLN. **a,** Flow cytometry analysis of fate- mapped C1 MLN cells from *RORγt-eGFP;mKate2-ON^Δzbtb46^* mice, gated for the indicated cell subsets (ILC3 were gated as TCR*β*^-^, TCR*γδ*^-^, B220^-^, MHCII^+^, ROR*γ*t-eGFP^+^, CCR6^+^, CD25^+^ and JC as TCR*β*^-^, TCR*γδ*^-^, B220^-^, MHCII^+^, ROR*γ*t-eGFP^+^, CD25^-^). Note that there is incomplete excision of the transcriptional stop signal by *zbtb46-Cre*. **b,** Representative image of cell-cell interactions of recently primed Hh-specific T cells with DC and ROR*γ*t-expressing cells. *Nur77- eGFP* tracer-labeled Hh7-2 T cells were transferred into of *RORγt-eGFP;mKate2-ON^zbtb46^* Hh- colonized mice. Cell colocalization of primed Hh7-2 (tracer dye^+^, GFP^+^) or naïve Hh7-2 (tracer dye^+^, GFP^-^) T cells with cDC (mKate2^+^ with dendritic morphology), ROR*γ*t-expressing cells (eGFP^+^, mKate2^+^ or eGFP^+^ alone with amoeboid morphology), or both were visualized using intravital multiphoton microscopy of the C1 MLN at 15 h after transfer. Note that Cell Tracer fluorescent labeling provides clear spatial discrimination of *RORγt-eGFP* and *Nur77-eGFP* expressing cells. **c,** Quantification and graphical representation of the total and individual rates of interaction of ROR*γ*t-expressing cells or cDC populations with primed or naïve Hh7-2 T cells. Data summarize cell-cell interactions from six 0.25mm^3^ three-dimensional regions of C1 MLN, (n=72 total Hh7-2 T cells), (n=49 primed and 23 naïve Hh7-2 T cells). All statistics were calculated by unpaired two-sided Welch’s t-test. Error bars denote mean ± s.d. *p*-values are indicated in the figure.

**Extended Data Fig. 12.**
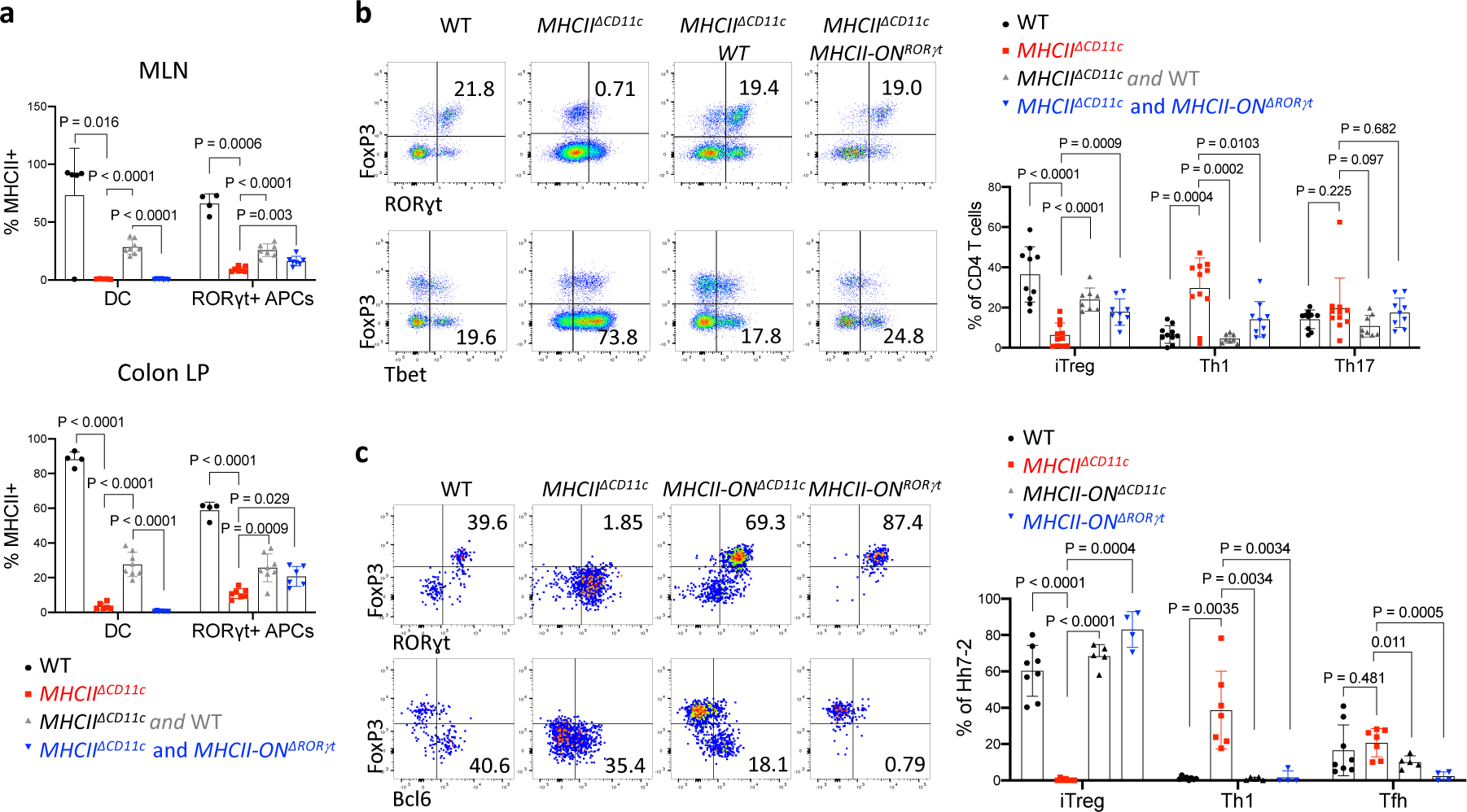
Gain-of-function expression of MHCII in ROR*γ*t^+^ cells rescues bone marrow-derived iTreg cell differentiation. **a,** Aggregate data showing MHCII frequency on donor-derived DC and ROR*γ*t^+^ cells in MLN and colon lamina propria from chimeric mice reconstituted with combinations of donor BM cells as indicated, with representative flow cytometry panel in Fig. 5b. MLN: WT (n=5), *MHCII^ΔCD11c^* (n=5), *MHCII^ΔCD11c^* and WT (n=8), *MHCII^ΔCD11c^* and *MHCII-ON^ΔRORγt^* (n=7). Colon: WT (n=4), *MHCII^ΔCD11c^* (n=6), *MHCII^ΔCD11c^* and WT (n=8), *MHCII^ΔCD11c^* and *MHCII-ON^ΔRORγt^* (n=6). **b,** Donor bone marrow-derived CD4^+^ T cell differentiation in colon lamina propria from chimeric mice reconstituted with combinations of BM cells as indicated. Representative flow panels (left) and aggregate data (right). WT (n=10), *MHCII^ΔCD11c^* (n=11), *MHCII^ΔCD11c^* and WT (n=8), *MHCII^ΔCD11c^* and *MHCII-ON^ΔRORγt^* (n=9). Colon: WT (n=4), *MHCII^ΔCD11c^* (n=6), *MHCII^ΔCD11c^* and WT (n=8), *MHCII^ΔCD11c^* and *MHCII-ON^ΔRORγt^* (n=7). **c,** Representative flow cytometry (left) and aggregate data (right) of Hh7-2 T cell differentiation in colon lamina propria of Hh-colonized bone marrow chimeric mice reconstituted with cells of indicated genotypes, 12 days after transfer of naive TCR transgenic T cells. WT (n=8), *MHCII^ΔCD11c^* (n=7), *MHCII-ON^ΔCD11c^* (n=5), and *MHCII-ON^ΔRORγt^* (n=4). Data summarize two or three independent experiments. All statistics were calculated by unpaired two-sided Welch’s t- test. Error bars denote mean ± s.d. *p-*values are indicated in the figure.

**Extended Data Fig. 13.**
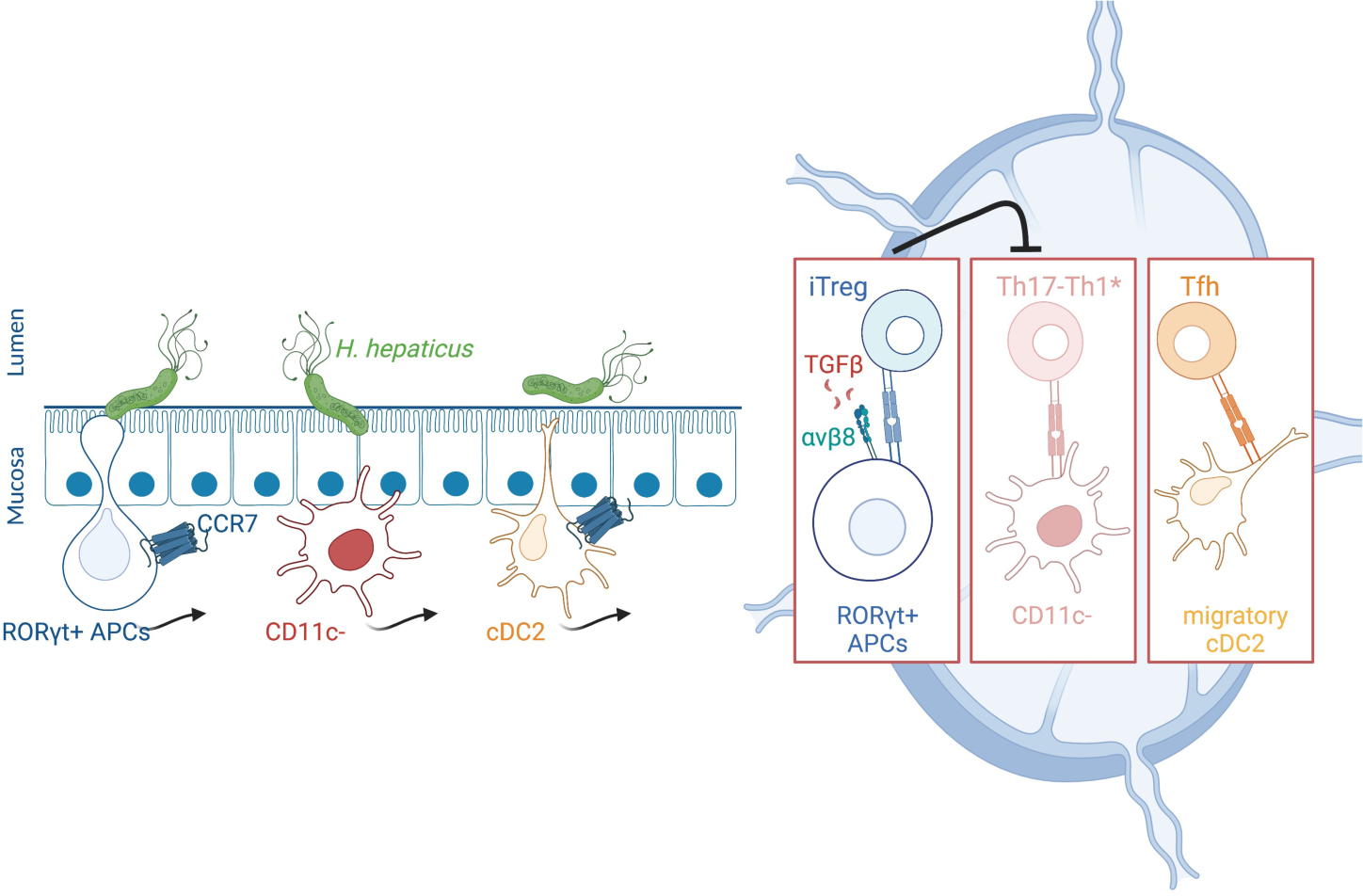
Schematic of the requirement of distinct APC subsets for T cell differentiation. CCR7 and integrin *α*vβ8 are required in ROR*γ*t^+^ APCs for iTreg cell differentiation. Note that other APCs, with differential requirements for CCR7 expression, are involved in the priming and differentiation of pathogenic Th17 and Tfh cells.

